# LncRNA *TAAL* is a Modulator of *Tie1*-Mediated Vascular Function in Diabetic Retinopathy

**DOI:** 10.1101/2024.09.13.612383

**Authors:** Gyan Ranjan, Samriddhi Arora, Sarmeela Sharma, Lakshita Sharma, Rahul C Bhoyar, Vigneshwar Senthivel, Vinod Scaria, Subhabrata Chakrabarti, Inderjeet Kaur, Sridhar Sivasubbu, Rajender K Motiani

## Abstract

Diabetic retinopathy (DR), a leading cause of vision impairment and blindness, is characterized by abnormal retinal vascular changes due to chronic hyperglycemia. The *Tie-1* signaling pathway, essential for vascular growth and remodeling, has emerged as a key therapeutic target, though its molecular mechanisms and interactome remain largely unclear. Through a protein-centric approach, we identified a novel lncRNA and named it *Tie1-associated angiogenic lncRNA (TAAL)*. *TAAL* lncRNA regulates endothelial cell migration, proliferation, tube formation, and permeability by modulating ER-calcium homeostasis and cytoskeleton dynamics. In zebrafish, *taal* modulation led to angiogenic defects, which were rescued by human *TAAL* orthologue. Our molecular studies further revealed that *TAAL* negatively regulates *Tie1* protein via ubiquitin-mediated degradation. Notably, *TAAL* expression is upregulated in the blood of DR patients and downregulated in endothelial DR cell models. Overexpression of *TAAL* restored endothelial permeability and VE-cadherin surface expression. These findings establish *TAAL* as a novel regulator of *Tie1* protein turnover, with potential therapeutic implications for diabetic retinopathy.

## Synopsis

*Tie1* is an endothelial protein that plays a critical role in neovascularization. Its interactome is not well studied. Here, we discover a novel lncRNA *TAAL* that interacts with *Tie1* protein and regulates its protein turnover thereby, *TAAL* modulates endothelial pathophysiology.

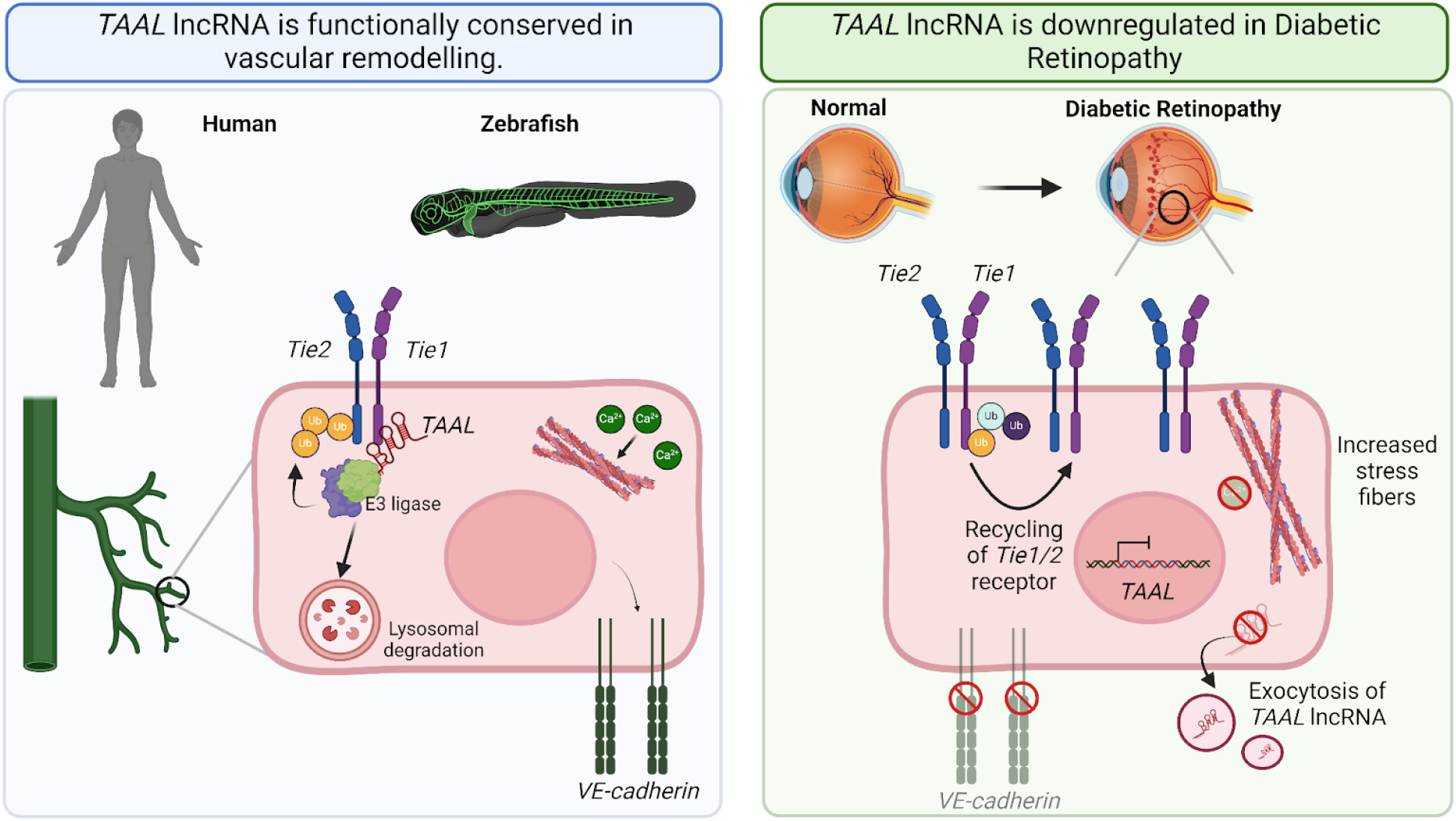

● The *TAAL* lncRNA interacts with the *Tie1* protein and regulates vascular remodeling.
● The *TAAL* lncRNA is functionally conserved between human and zebrafish. It plays an important role in regulating vascular angiogenesis during development.
● The *TAAL* lncRNA controls the protein turnover of *Tie1* by ubiquitin-mediated modulation of its stabilization and translation.
● The *TAAL* lncRNA is hyperactive during diabetic retinopathy and is a potential biomarker and a potent therapeutic target.

## Introduction

Diabetes mellitus (DM) is a chronic metabolic disorder characterized by persistent hyperglycemia, which can have detrimental effects on multiple organs, including the kidneys, liver, and cardiovascular system. Among the most serious complications of DM is diabetic retinopathy (DR), a condition driven by abnormal retinal neovascularization and dysregulated angiogenesis. These pathological changes increase vascular permeability, leading to retinal hemorrhage and, in severe cases, vision impairment or blindness (Wong *et al*, 2016; Duh *et al*, 2017). It is estimated that approximately 1 in 5 individuals with diabetes develop some degree of DR, with the incidence being particularly high in low- and middle-income countries where late diagnosis is common due to resource constraints. Early detection of DR is crucial, as timely intervention can prevent up to 95% of vision impairment and blindness. Current treatment strategies include laser therapy, vitrectomy, and anti-VEGF therapies (Wong *et al*, 2016; Duh *et al*, 2017). However, the prevalence of DR is rising rapidly, and there remains a significant unmet need for cost-effective, point-of-care diagnostic tools and targeted second-generation therapies to manage better and control the progression of this condition. Recent studies have shown that angiopoietin and *Tie* signaling plays a crucial role in protecting vascular dysfunction. During hyperglycemic conditions, the Ang-Tie signaling is severely affected (Saharinen *et al*, 2017).

*Tie1* is a major effector protein during hyperglycemic conditions as it is a major player in neovascularization. Although its interactome and ligand remain a mystery, and it is often termed an orphan receptor (Mueller & Kontos, 2016; Sato *et al*, 1995). Recent investigations have shed light on their more complex involvement in regulating endothelial homeostasis and remodeling. *Tie1* in zebrafish and mice has been shown to affect endothelial cell angiogenesis and proper homeostasis *in vivo* (Carlantoni *et al*, 2021; Reinardy *et al*, 2015; D’Amico *et al*, 2010).

Moreover, functional studies indicate that *Tie1* plays a dual role in angiogenesis: it exerts a negative regulatory effect on *Tie2* levels within angiogenic tip cells, while positively regulating *Tie2* by forming heterodimers in stalk cells (Savant *et al*, 2015). These heterodimers are crucial for the maturation and remodeling of vessels through angiopoietin 1 signaling. Previous investigations suggested that Angiopoietin-1 binding independently to Tie2 and in heterodimeric association with *Tie1*, activating downstream signaling kinases such as PI3K/Akt pathway or Rho/Rac1, thereby modulating endothelial cell proliferation, migration, and survival(Saharinen *et al*, 2017; Joussen *et al*, 2021; Papapetropoulos *et al*, 2000; Saharinen *et al*, 2008). The regulation of *Tie1* and *Tie2* levels in vascular remodeling has been a focal point of research due to their potential as targeted therapies. Their involvement in diseases such as cancer metastasis and diabetic retinopathy underscores their clinical significance (Saharinen *et al*, 2017; Huang *et al*, 2010; Joussen *et al*, 2021; Augustin *et al*, 2009). It has prompted intensive investigation as they are the key player in angiogenesis and endothelial function.

Long noncoding RNAs (lncRNAs) are a significant and abundant class of regulatory molecules in endothelial cells, where they play a crucial role in controlling gene expression at various levels, including chromatin remodeling, transcription, and post-transcriptional processing. Recent discoveries, such as lncRNAs like *VEAL2, GATA-AS, SENCR, LASSI, STEEL, MANTIS, LEENE,* and *Tie1-AS*, have underscored their essential functions in regulating endothelial cell proliferation, migration, permeability, and stability. These lncRNAs have been extensively studied, revealing new mechanisms of action and interactions that continue to emerge. In addition to their regulatory roles, lncRNAs have shown promise as biomarkers for vascular diseases, given their high specificity, offering potential for early diagnosis and personalized treatment strategies. Given their pivotal role in vascular biology, lncRNAs are increasingly recognized as both therapeutic targets and biomarkers for managing conditions like diabetic retinopathy, where novel RNA-based interventions could provide significant clinical benefits.

This study explores the RNA interactions of *Tie1*, a key protein involved in neovascularization and complications in diabetic retinopathy (DR). Through a Tie1 pulldown approach, we identified a previously uncharacterized and conserved lncRNA, *TAAL*, in human endothelial cells and zebrafish. Our research highlights the critical role of lncRNA *TAAL* in fundamental physiological processes such as tube formation and vascular permeability in endothelial cells, with significant implications for vascular development in zebrafish embryos. Notably, we discovered a crucial interaction between lncRNA *TAAL* and the *Tie1* protein, where *TAAL* is essential for modulating *Tie1* protein stability. This relationship was further examined in the context of DR, where lncRNA *TAAL* levels were notably upregulated in patient samples compared to controls. These findings emphasize the importance of lncRNA *TAAL* in regulating *Tie1-*mediated angiogenesis, permeability and vascular functions, particularly under diabetic retinopathy conditions.

## Results

### Protein-centric discovery of novel lncRNA - *Tie1*-associated angiogenic lncRNA (*TAAL*)

*Tie-1* plays a critical role in vascular development, as demonstrated in the mouse and zebrafish models(Cao *et al*, 2023; Savant *et al*, 2015; Carlantoni *et al*, 2021). However, *Tie-1*’s interactome and the molecular mechanisms regulating its expression remain poorly understood. Here, we performed a protein-targeted pulldown of *Tie1* and determined if any *Tie1* protein-associated long non-coding RNAs are involved in the angiogenesis pathway using HUVEC-hTert2 cell line.

We performed RNA immunoprecipitation (RIP) of *Tie1* protein followed by RNA-sequencing (Figure 1A). After applying a stringent cutoff filter of greater than 5 fold change and -log_10_ (P-value) value greater than 2, we identified a bi-exonic polyadenylated lncRNA (Supply Fig 1A) and named it *TIE1-associated angiogenic lncRNA (TAAL) (previous names- ENST00000457948.3, NONHSAG007003.2, RP11-79M19.2)(*Figure 1B). The lncRNA originates from the intron between 2 and 3 exons of the GRK5 gene (Figure 1C). Further, it does not exhibit any coding potential as computed using bioinformatic algorithms (Coding Potential Assessment Tool (CPAT2) coding probability - 0.0003 (Wang *et al*, 2013); Coding Potential Calculator (CPC2) coding probability - 0.0216282 (Kang *et al*, 2017))(Supply Fig 1B) . We also performed ribosomal pulldown followed by RT-qPCR of the *TAAL* lncRNA and did not find any ribosome enrichment on lncRNA *TAAL*, further suggesting it to be non-coding (Supply Fig 1C). The second exon and promoter of lncRNA *TAAL* is observed to be conserved in mammals and birds but is lost in reptiles and fish, exhibiting no sequence conservation. It also displays all the epigenomic marks, including H3K27ac, H3K4me1, H3K4me3, and CAGE reads from ENCODE data in HUVECs, suggesting it is a genuine transcript (Figure 1C) (ENCODE Project Consortium *et al*, 2020). Expression profiling of the lncRNA using the GTEx v8 portal exhibited it highly expressed in the testis, ovary, pituitary, lungs, and heart tissues (Supply Fig 2A)(Ferraro *et al*, 2020). Next, we performed RIP-RT-qPCR to determine if the *TAAL* lncRNA also interacts with *Tie2*. We did not find any enrichment of the lncRNA to the *Tie2* protein (Figure 1D). Furthermore, to understand its localization, we performed subcellular fractionation and qRT-PCR, which suggested that the *TAAL* lncRNA is cytoplasmic enriched (Figure 1E). Single-molecule RNA *FISH* (*smFISH*) also demonstrates that *TAAL* lncRNA is enriched in the cytoplasm (Supply Fig2B). We also performed co-immunofluorescent and *smFISH* for *TAAL* lncRNA, *Tie1*, and *Tie2* proteins and observed colocalization of the lncRNA and proteins (Figure 1F). The interaction of *TAAL* lncRNA along with the *Tie1* protein suggests its *trans*-role in regulating protein function.

**Figure 1:**
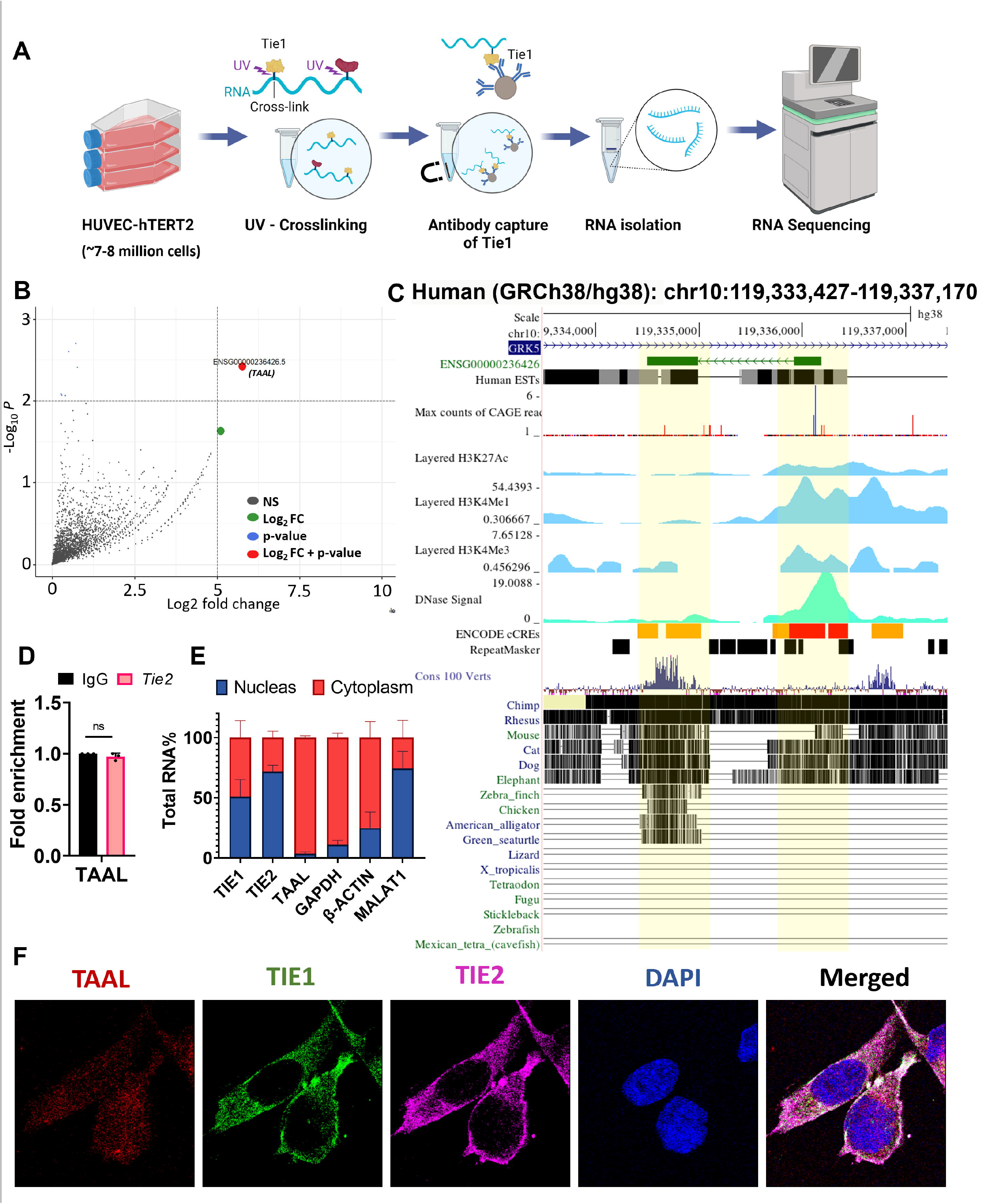
Tie1-associated angiogenic lncRNA (TAAL) interaction Tie1 protein. [A] Schematic of RNA-immunoprecipitation pipeline for Tie1 protein using HUVEC-hTert2 cell line. [B] Differential expression analysis of *Tie1/IgG* revealed one enriched lncRNA with a fold change greater than 5, represented in red. [C] UCSC genome browser snapshot of *TAAL lncRNA* locus in human. [D] The bar plot represents RNA-immunoprecipitation of *Tie2* in HUVEC-hTert2 cells and shows no enrichment of lncRNA *TAAL.* Data from 3 independent biological replicates plotted as mean fold change ± standard deviation . ns- not significant (two-tailed unpaired t-test) [E] The bar plot depicts the relative abundance of Tie1, Tie2, *TAAL*, GAPDH, β-actin, and MALAT1 in various subcellular fractions quantified using qRT-PCR. *TAAL*, GAPDH, and β-actin show enrichment in the cytoplasmic fraction. Tie1 and Tie2 exhibit equal enrichment in both the cytoplasm and nucleus, while *MALAT1* is predominantly enriched in the nucleus. The data represent relative abundance percentages ± SEM from three independent experiments. [F] Single molecular FISH (smFISH) of *TAAL* (610nM) and Co-IF for Tie1 (GFP) and Tie2 (Magenta) highlight their colocalization. Magnification-60x

### LncRNA TAAL modulates the pathophysiological functions of endothelial cells

To understand the functionality of the lncRNA, we performed siRNA-mediated knockdown using siTAAL and overexpression using pcDNA3.1-TAAL plasmid. We observed significant knockdown and overexpression of the *TAAL* lncRNA transcript *in vivo* in the HUVEC-hTert2 cell line (Supply Fig 3 A-B). Next, we conducted scratch-wound assays and transwell migration assays to elucidate the migratory properties of cells upon lncRNA *TAAL* manipulation. Remarkably, the knockdown of lncRNA *TAAL* promoted enhanced wound closure and increased the migratory capacity of cells. In contrast, overexpression of lncRNA *TAAL* exerted a decrease in wound closure rate and an inhibitory effect on cell migration (Figure 2A-D). Additionally, BrdU proliferation assays corroborated our findings, demonstrating increased cell proliferation following lncRNA *TAAL* knockdown, while overexpression of lncRNA *TAAL* attenuated cellular proliferation (Figure 2E-F). Moreover, cell cycle analysis unveiled distinct alterations in cell cycle distribution patterns associated with lncRNA *TAAL* modulation. Specifically, the knockdown of lncRNA TAAL led to an accumulation of cells in the G0/G1 phase, indicative of an active proliferative state. In contrast, overexpression of lncRNA TAAL showed no significant change (Supply Fig I-J). Furthermore, we conducted a permeability assay using a collagen-coated transwell plate to assess the impact of lncRNA *TAAL* on cellular permeability. We observed a significant increase in permeability following the knockdown of lncRNA *TAAL* (Figure 2G). Conversely, we noted a decrease in permeability upon the overexpression of lncRNA *TAAL*, suggesting lncRNA’s role in maintaining cellular barrier integrity (Figure 2H).

**Figure 2:**
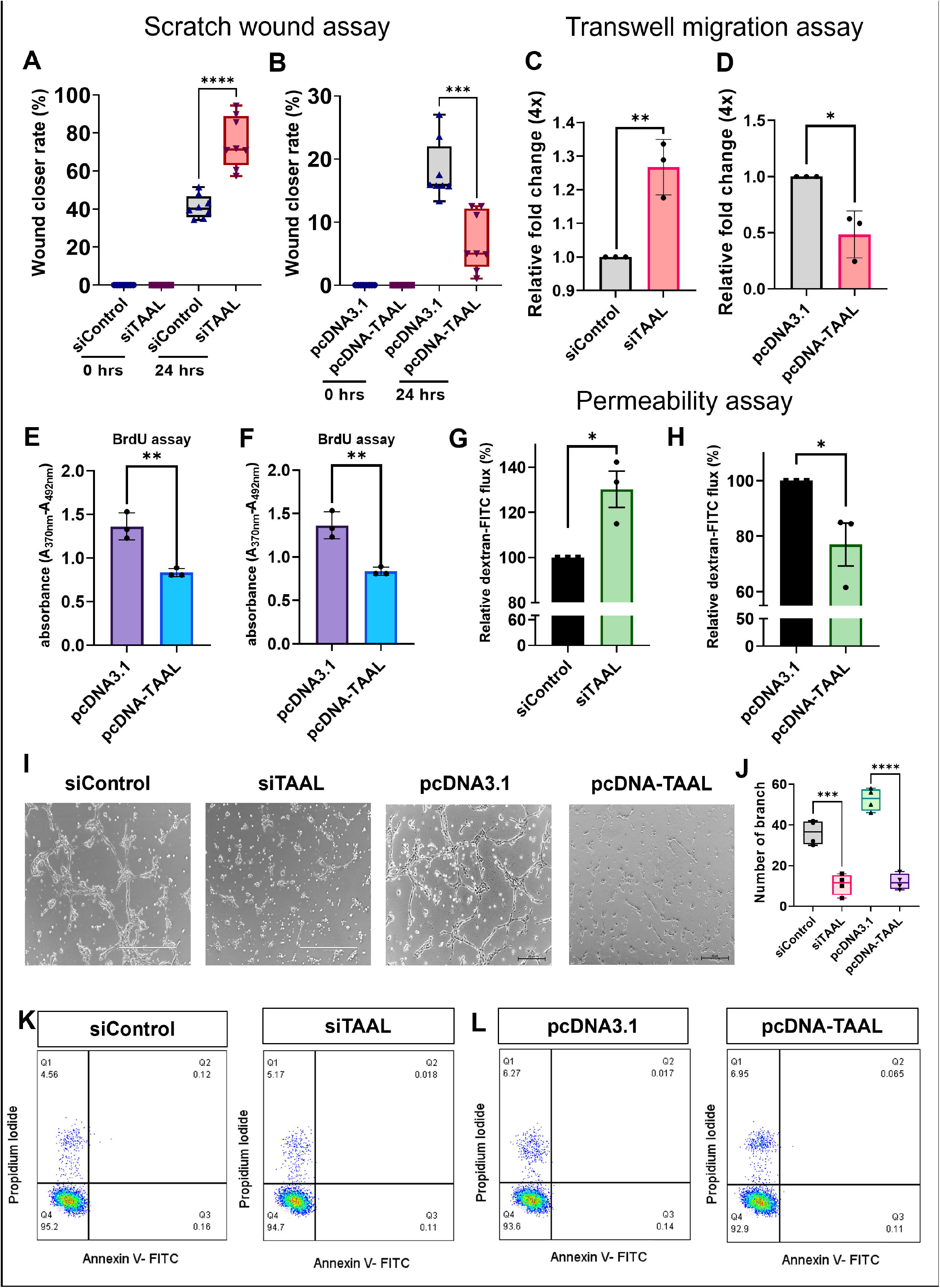
lncRNA *TAAL* modulates pathophysiological function in endothelial cells. [A-B] Box and whisker plot representing quantification of wound closer rate in scratch wound assay after 24 hrs in control and *TAAL* knockdown or overexpressed cells. *** p-value ≤ 0.001 ; **** p-value ≤ 0.0001. (two-tailed unpaired t-test). [C-D] Bar plot representing quantification of cells migration in the transwell assay compared in control and *TAAL* knockdown or overexpressed cells.* p-value ≤ 0.05; ** p-value ≤ 0.01 (two-tailed unpaired t-test). [E-F] Bar plot representing quantification of absorbance in the BrdU proliferation assay compared in control and *TAAL* knockdown or overexpressed cells. ** p-value ≤ 0.01 (two-tailed unpaired t-test). [G-H]Knockdown and overexpression of *TAAL* significantly alter the permeability of dextran-conjugated FITC. Bar plot representing relative dextran-FITC efflux in control and *TAAL* knockdown or overexpressed cells. * p-value ≤ 0.05. (two-tailed unpaired t-test). [I]Knockdown and overexpression of *TAAL* in HUVECs exhibited a reduction in tube formation in Matrigel when compared to the control. [J] Box and whisker plot representing quantification of the number of branched vessels in *TAAL* knockdown (siTAAL) and overexpressed (pcDNA-TAAL) cells along with their controls (siControl; pcDNA3.1). *** p-value ≤ 0.001, **** p-value ≤ 0.0001. (two-tailed unpaired t-test). [K-L] Flow cytometry analysis of cell viability, stained by Annexin V (Q3) and PI (Q1) with viable cells which are negative for both Annexin V and PI (Q4) in control and *TAAL* lncRNA knockdown or overexpression cells.

Next, we evaluated the angiogenic tube-forming function of the HUVEC-hTert2 cell under knockdown and overexpression conditions. We observed that lncRNA *TAAL* knockdown cells *(siTAAL)* were forming colonies but could not attain the tubular morphology on matrigel compared to control transfection *(siControl)*. Similarly, upon *TAAL* lncRNA overexpression (*pcDNA-TAAL*), we observed that cells could not attain the tubular morphology as seen in the control pcDNA-3.1 transfection condition (Figure 2I-J). Next, we investigated cell viability’s potential influence on the perturbations in tubulogenesis observed upon manipulating lncRNA *TAAL* expression levels. To comprehensively assess cell viability, we performed Annexin-V/PI staining assays. We observed no significant change in cell viability following transfection with siTAAL and pcDNA-*TAAL* constructs, suggesting the effects are not due to cellular toxicity but rather the modulation of lncRNA *TAAL* expression (Figure 2K-L). These data together suggest that lncRNA *TAAL* modulates the physiological function of endothelial cells.

### LncRNA TAAL regulates calcium-dependent cytoskeletal and junctional dynamics

Given the observed alterations in proliferation and migration, we investigated the impact of lncRNA *TAAL* modulation on cellular cytoskeletal dynamics. In the endothelial cells, we stained the F-actin using Alexa Fluor 488 phalloidin to visualize cytoskeletal structures. Our analysis revealed noteworthy changes in cytoskeletal organization upon knockdown and overexpression of *TAAL* lncRNA. Specifically, *TAAL* lncRNA knockdown cells exhibited a marked increase in the length and density of stress fibers, indicative of enhanced cytoskeletal stiffness (Figure 3A).

**Figure 3:**
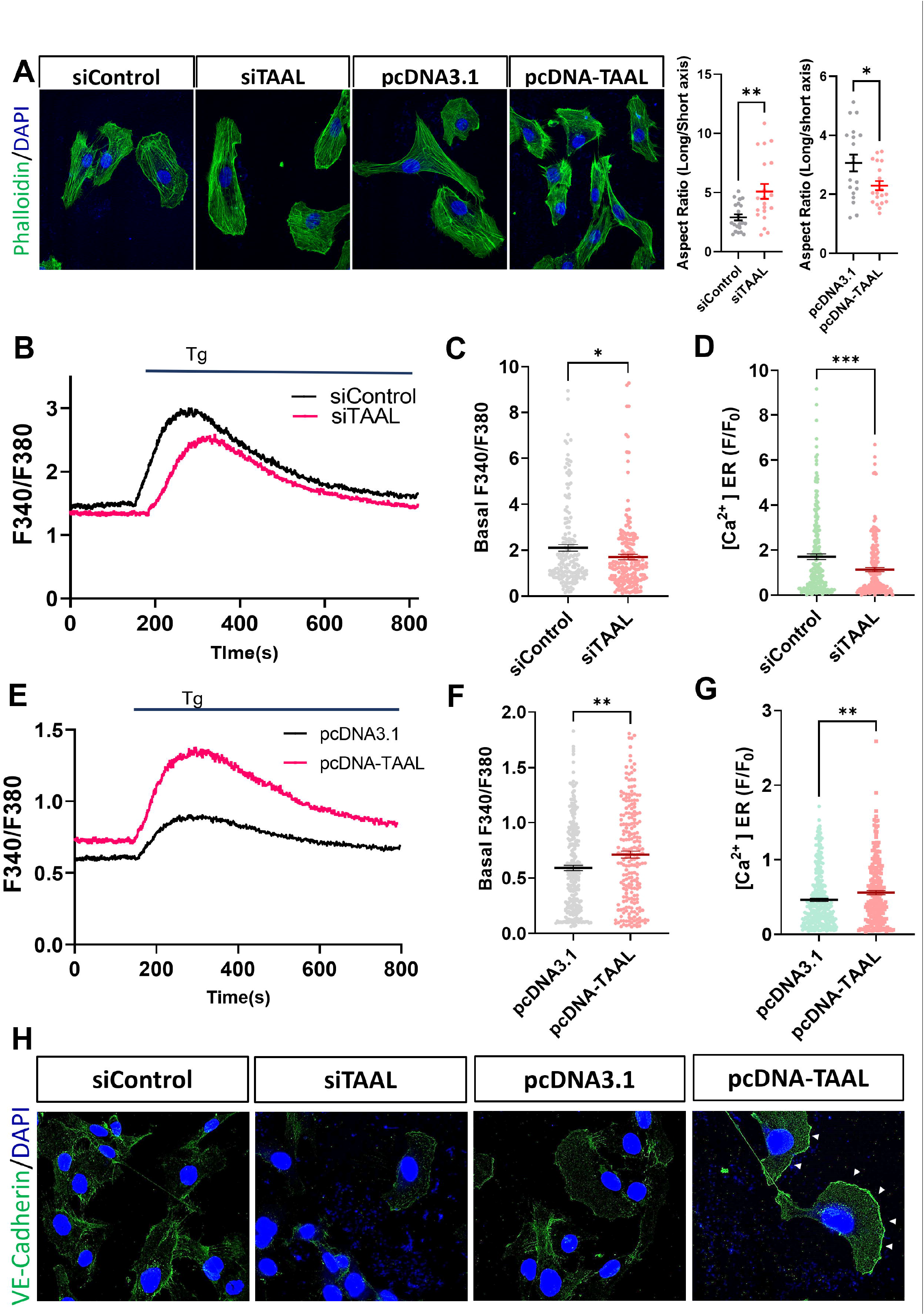
LncRNA *TAAL* modulates cytoskeleton dynamics by altering calcium levels. [A] Knockdown and overexpression of *TAAL* significantly alter the cytoskeleton dynamics in HUVEC cells. Confocal images of F-actin stained with Alexa Fluor 488 phalloidin in control and *TAAL* knockdown or overexpressed cells (60X magnification). Dot plot representing quantification of aspect ratio (long/short axis) in control and *TAAL* knockdown or overexpressed cells.* p-value ≤ 0.05; ** p-value ≤ 0.01 (two-tailed unpaired t-test). [B] Representation of Ca^2+^ imaging trace of control and *TAAL* knockdown HUVEC cells with stimulation with thapsigargin (Tg). [C] Quantification of basal Ca^2+^ levels in control and *TAAL* knockdown HUVEC cells. * p-value ≤ 0.05. (two-tailed unpaired t-test). [D] Quantification of Ca^2+^ levels release from the endoplasmic reticulum (ER) upon stimulation in Tg in control and *TAAL* knockdown HUVEC cells. *** p-value ≤ 0.001. (two-tailed unpaired t-test). [E] Representation of Ca^2+^ imaging trace of control and *TAAL* overexpressed HUVEC cells with stimulation with Tg. [F] Quantification of basal Ca^2+^ levels in control and *TAAL* overexpressed HUVEC cells. ** p-value ≤ 0.01. (two-tailed unpaired t-test). [G] Quantification of Ca^2+^ levels release from ER upon stimulation in Tg in control and *TAAL* overexpressed HUVEC cells. ** p-value ≤ 0.01. (two-tailed unpaired t-test). [H] Knockdown and overexpression of *TAAL* significantly alters the localization of VE-Cadhaerin at the junction of HUVEC cells. Confocal images of immunofluorescence VE-Cadhaerin in control and *TAAL* knockdown or overexpressed cells (60X magnification).

Furthermore, these cells elongated in cell length compared to their respective controls (Supply Fig 5A). Conversely, overexpression of *TAAL* lncRNA elicited contrasting effects, with cells displaying a reduction in both stress fiber length and cell size (Figure 3A). This suggests a role for *TAAL* lncRNA in modulating cytoskeletal dynamics and cellular stiffness, potentially influencing cellular morphology and migration ability.

Previous investigations have highlighted the involvement of endoplasmic reticulum (ER) calcium dynamics in mediating cytoskeletal remodeling in endothelial cells (Shinde *et al*, 2013; Tsai & Meyer, 2012; Tsai *et al*, 2014). Angiopoietin 1/2 mediated calcium regulation via the *Tie* receptors has also been previously investigated and has been linked to angiogenesis (Pafumi *et al*, 2015). Hence we investigated the impact of lncRNA *TAAL* modulation on intracellular calcium levels, specifically focusing on calcium regulation within the ER. We assessed changes in calcium dynamics by using live cell Ca^2+^ imaging with the ratiometric dye Fura2-AM. Additionally, we employed the pharmacological inhibitor Thapsigargin (Tg) to deplete ER Ca^2+^ stores, aiming to elucidate the role of *TAAL* in calcium homeostasis. Our observations revealed significant alterations in calcium homeostasis upon manipulating lncRNA *TAAL* expression levels. The knockdown of lncRNA *TAAL* resulted in a significant decrease in basal intracellular calcium levels and a reduction in ER calcium content (Figure 3B-D). Conversely, overexpression of lncRNA *TAAL* led to increased basal intracellular calcium levels and ER calcium content (Figure 3E-G). Consistent with previous findings in cell culture and zebrafish models, which have demonstrated that calcium spikes induce stress fiber contraction (Stolz & Bereiter-Hahn, 1988; Tiruppathi *et al*, 2002; Shinde *et al*, 2013; Dalal *et al*, 2020; Sahu *et al*, 2017), our results suggest that changes in cytoskeletal dynamics associated with lncRNA *TAAL* modulation may be mediated through alterations in calcium influx. We also checked the expression levels of a few calcium modulator genes and observed that STIM1 levels were downregulated in *TAAL* overexpressed cells (Supply Fig 6A-B). Also, upon modulation of lncRNA *TAAL*, the levels of *STIM2* were significantly altered; in the knockdown state, it was suppressed, and in overexpression of lncRNA, it was upregulated, correlating with the calcium levels inside the cell upon similar conditions (Supply Fig 6C-D). We also observed decreased ORAI2 leaves upon *TAAL* overexpression in the cells (Supply Fig 6 E-F). This suggests that intracellular calcium dynamics, upon modulation of lncRNA *TAAL*, are governed by alterations in the levels of *STIM* proteins,, a critical regulator of cellular and ER calcium homeostasis.

To further elucidate the underlying mechanisms contributing to changes in cellular permeability following lncRNA *TAAL* alteration, we assessed mRNA levels of *VEGFR2* (KDR) and VE-Cadherin (Hamdollah Zadeh *et al*, 2008; Phoenix *et al*, 2022; Bates & Curry, 1997; Dalal *et al*, 2020; Gavard, 2013; Sandoval *et al*, 2001) but observed no significant changes (Supplementary Fig 7A-D) . Next, we examined the localization of VE-Cadherin, an adhesion protein regulated by the *Tie1-Tie2* signaling pathway and calcium influx. Our analysis revealed that *TAAL* knockdown resulted in decreased membrane localization of VE-Cadherin, while *TAAL* overexpression increased its membrane localization (Figure 3H). These findings collectively suggest that modulation of lncRNA *TAAL* influences the VE-Cadherin membrane localization.

### TAAL lncRNA is functionally conserved in zebrafish and modulates angiogenesis

To check if the lncRNA is functionally conserved across species, we performed a protein-centric pulldown of *tie1* in zebrafish at 24 hpf, followed by RNA sequencing. Upon analysis, we observed a lncRNA *ZFLNCG09997* (hereafter called lncRNA *taal*) was 4.7 fold enriched and had a -log_10_ P-value greater than 15 (Figure 4A-B). The zebrafish *taal* lncRNA gene produces a 3 exonic polyadenylated lncRNA (Supply Fig 8A) with a transcript length of 2632bp observed on chromosome 18 of zebrafish, arises antisense 5’ upstream of endonuclease, polyU-specific C (*endouc*). The *taal* lncRNA is noncoding in nature, as assessed through the online tool Coding Potential Calculator (CPC2) (coding probability -0.0403034), and has no potential predicted ORF. The *taal* lncRNA shows no sequence conservation with its human orthologue *TAAL*.

**Figure 4:**
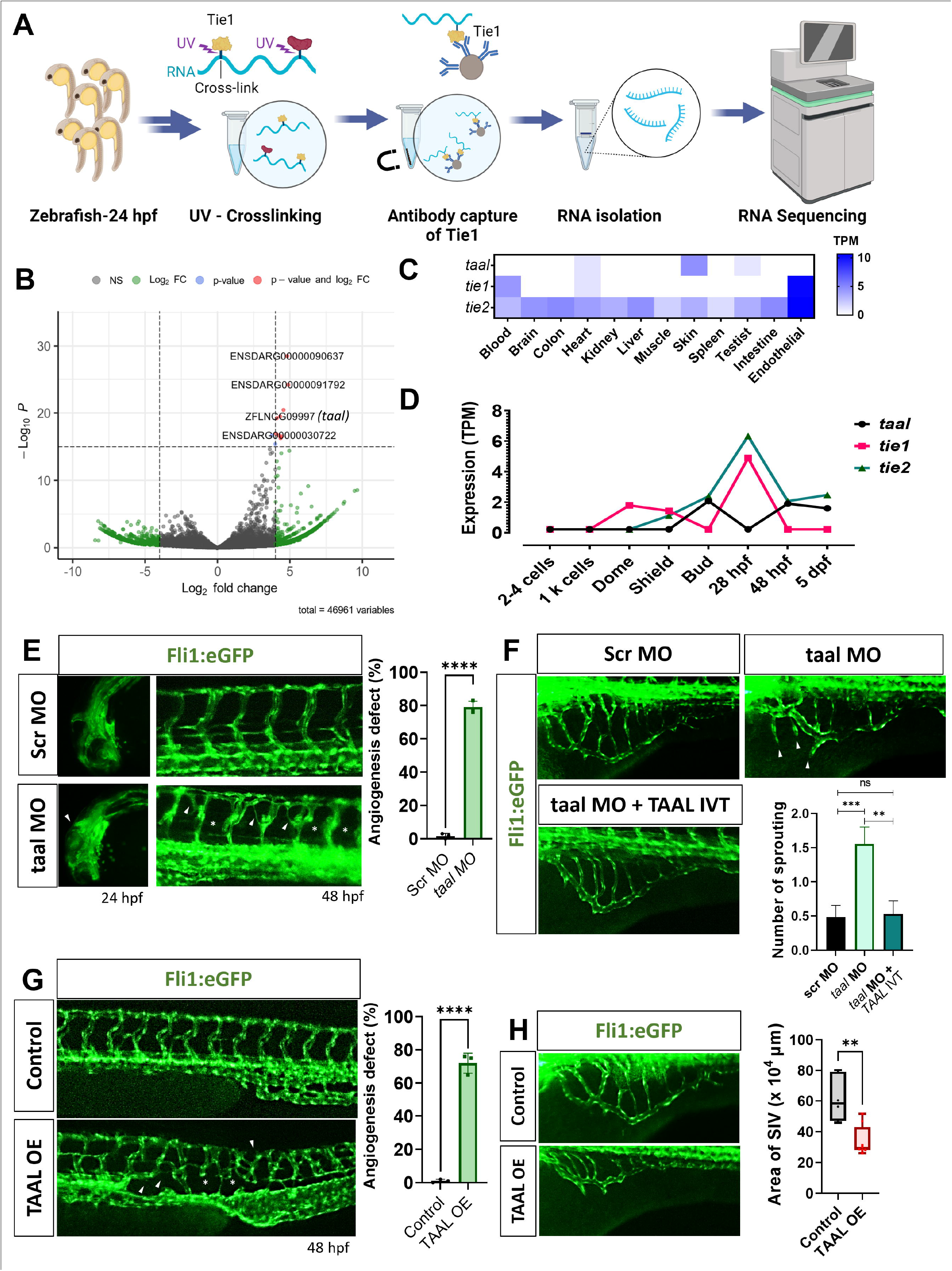
Conserved function of lncRNA *TAAL* in zebrafish regulates angiogenesis. [A] Schematic of RNA-immunoprecipitation pipeline for *tie1* protein using24 hpf zebrfish embryos. [B] Differential expression analysis of tie1/IgG revealed one enriched lncRNA (ZFLNCG0999) with a fold change greater than 4.5 and log_10_ P value greater thant 15, represented in red. [C] Expression pattern of *taal* lncRNA, *tie1* and *tie2* (measured in TPM) across eight distinct developmental stages in zebrafish, utilizing publicly accessible RNA-seq data. [D] Expression pattern of *taal* lncRNA, *tie1* and *tie2* (measured in TPM) across 12 different tissues using publically available RNA-seq data. [E] Representative images of cranial and trunk vasculature of control and morpholino mediated *taal* knockdown Tg (fli1:eGFP) zebrafish at 24 hpf and 48 hpf at 5x magnification. Bar graph represent quantification of angiogenic defects in control and *taal* knockdown zebrafish. Arrow marks represent the defective angiogenic vessel. **** p-value ≤ 0.0001. (two-tailed unpaired t-test). [F] LncRNA *taal* knockdown enhances angiogesis in zebafish. Representative images of angiogenic sprouting in control and morpholino *taal* injected at at subintestinal vessels (SIVs) in Tg (fli1:eGFP) zebrafish. Complementation of human lncRNA *TAAL* in morpholino-*taal* injected Tg(fli1:eGFP) zebrafish at subintestinal vessels (SIVs) halted angiogenic sprouting. Arrow marks represent the sprouting vessels. Bar graph represent quantification of number of angiogenic sprotingin in control, *taal* knockdown and *taal* knockdown complimented with human *TAAL* in zebrafish. ns- not significant; ** p-value ≤ 0.01; *** p-value ≤ 0.001. (two-tailed unpaired t-test). [G] Representative images of trunk vasculature of control and *TAAL* overexpression (OE) in Tg (fli1:eGFP) zebrafish at 48 hpf with 5x magnification. Bar graph represent quantification of angiogenic defects in control and *TAAL* overexpressed zebrafish. Arrow marks represent the defective angiogenic vessel. **** p-value ≤ 0.0001. (two-tailed unpaired t-test). [F] Representative images of angiogenic sprouting in control and Overexpressed *TAAL* injected at at subintestinal vessels (SIVs) in Tg (fli1:eGFP) zebrafish. Box and whisker plot repent the area of SIVs in control and Overexpressed *TAAL* injected zebrafish. ** p-value ≤ 0.01 (two-tailed unpaired t-test).

The expression profile analysis across adult tissues and developmental stages using publicly available RNA seq data (Pauli *et al*, 2012; Yang *et al*, 2020; Sehgal *et al*, 2021) unveiled the expression of the lncRNA in zebrafish’s heart, testis, and skin (Figure 4C). We also observed a prominent expression peak of the lncRNA *taal* during the bud stage, followed by sustained expression at 48 hours post-fertilization (hpf) and 5 days post-fertilization (dpf) (Figure 4D). Intriguingly, an inverse expression pattern was observed for the protein-coding genes *tie1* and *tie2* at 24 hpf, where their expression levels were markedly elevated while lncRNA *taal* expression remained minimal. Notably, the initiation of angiogenesis from the posterior cardinal vein (PCV) occurs during this developmental stage, implicating *tie1* and *tie2* in the sprouting and angiogenesis process. The observed expression pattern of lncRNA *taal* suggests a potential inhibitory role for *tie1* and *tie2* in angiogenesis during this critical developmental stage.

An antisense splice-blocking morpholino-based knockdown approach was employed to elucidate further the functional significance of lncRNA *taal* in zebrafish development. Knockdown of lncRNA *taal* resulted in notable defects in zebrafish angiogenesis, particularly evident in patterning defects observed in the intersegmental vessels (ISVs) at 2 dpf (Figure 4E). Additionally, investigation into the role of lncRNA *taal* in angiogenesis revealed enhanced angiogenic sprouting from the subintestinal vessels (SIVs) in zebrafish injected with taal morpholino near the SIV site as compared to control injections with scramble morpholino, indicative of an anti-angiogenic role for lncRNA *taal*. Further confirmation of lncRNA *taal’s* functional conservation across species was attained through complementation and rescue assays. Injection of a wild-type copy of human lncRNA *TAAL* demonstrated a reduction in angiogenic sprouting when co-injected with *taal* morpholino, affirming the orthologous nature of lncRNA *taal* between humans and zebrafish (Figure 4F). Moreover, an overexpression experiment of lncRNA *TAAL* injected into the single-cell zebrafish embryos revealed abnormal vascular patterning (Figure 4G). Furthermore, the injection of *TAAL* lncRNA at the SIV site led to a reduction in the area of the SIV vessel formation (Figure H). This suggests a pivotal role of zebrafish *taal* lncRNA in modulating angiogenic processes. These findings provide compelling evidence that zebrafish *taal* lncRNA is a functional orthologue to human *TAAL* lncRNA and has an important role in modulating and controlling angiogenic function in vertebrates.

### LncRNA TAAL negatively regulates Tie1/2 protein turnover

To understand the molecular mechanism we examined the expression pattern of *Tie1* and *Tie2* protein, with which the lncRNA *TAAL* interacts. We observed an increase in the protein expression upon knockdown of the lncRNA *TAAL* and a decrease in the expression upon overexpression of the lncRNA (Figure 5A). We also estimated *Tie1* and *Tie2* mRNA levels and found no significant change in their mRNA expression upon knockdown and overexpression of the lncRNA *TAAL* (Figure 5B). These findings indicate that lncRNA *TAAL* post-translationally regulates the levels of *Tie1* and *Tie2* in endothelial cells. To further investigate this regulation, we conducted immunofluorescence analysis of *Tie1* and *Tie2* under lncRNA *TAAL* knockdown and overexpression conditions. In *TAAL* knockdown cells, we observed increased *Tie1* and *Tie2* protein expression. Additionally, there was increased clustering of *Tie1* proteins and enhanced colocalization of *Tie1* and *Tie2* within the cells. Conversely, overexpression of lncRNA *TAAL* significantly reduced the *Tie1* and *Tie2* protein signals and decreased colocalization of *Tie1* and *Tie2* (Figure 5C).

**Figure 5:**
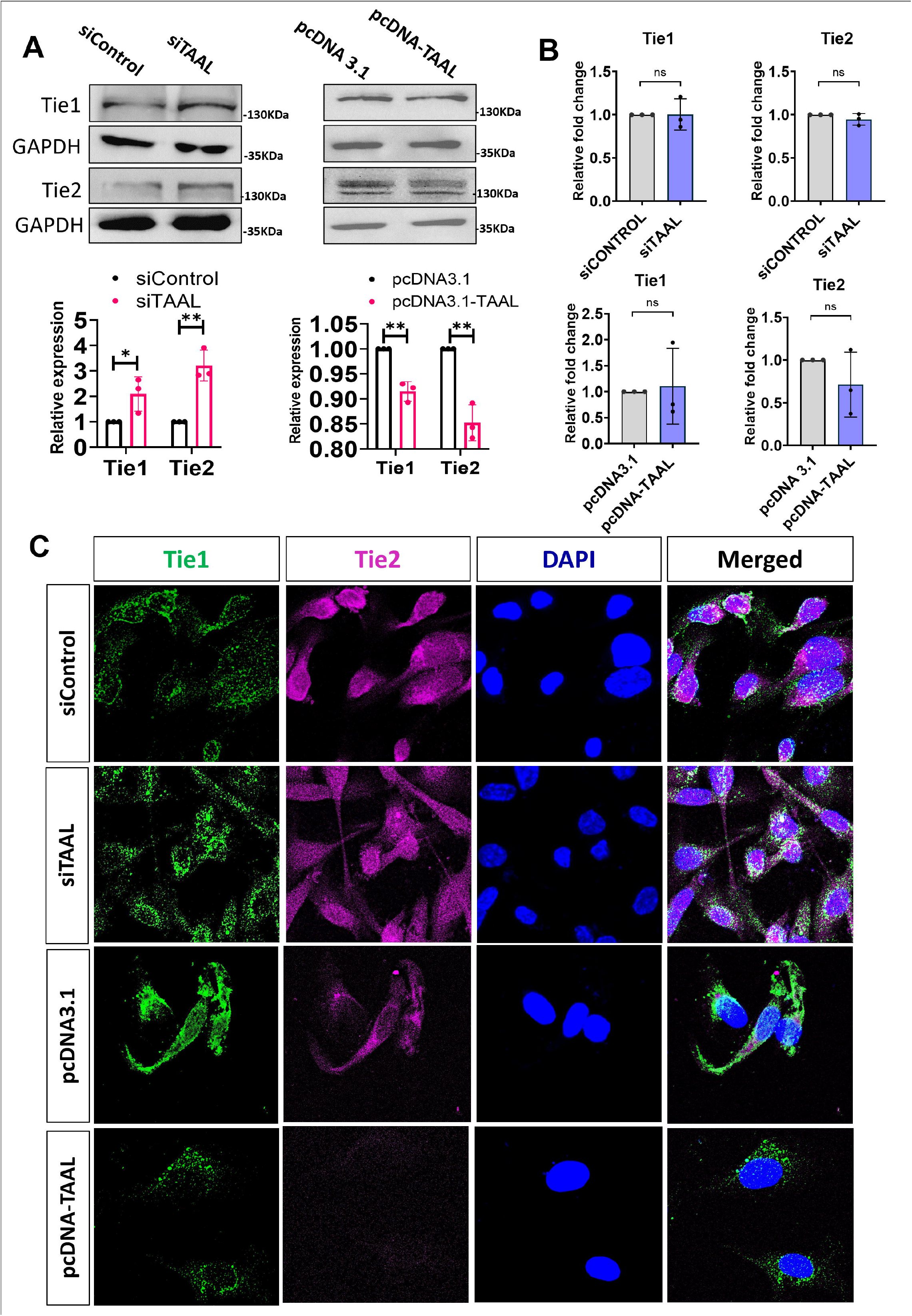
LncRNA *TAAL* modulates *Tie1* and *Tie2* protein levels. [A] Immunoblotting of Tie1 and Tie2 in control and TAAL knockdown or overexpressed cells. Bar plot representing quantification of blots. * p-value ≤ 0.05; ** p-value ≤ 0.01 (two-tailed unpaired t-test). [B] Bar plot representing relative fold change in mRNA expression levels of *Tie1* and *Tie2* between control and *TAAL* knockdown or overexpressed cells. ns- not significant (two-tailed unpaired t-test). [C] Knockdown and overexpression of *TAAL* significantly alters the protein levels of Tie1 and Tie2 in HUVEC cells. Confocal images of immunofluorescence Tie1 and Tie2 in control and *TAAL* knockdown or overexpressed cells (60X maginication).

### LncRNA TAAL modulates Tie1/2 protein turnover by ubiquitination

To elucidate the mechanisms underlying *Tie1/2* protein turnover, we employed proteomics using LC/MS mass spectrometry on *TAAL* knockdown cells to identify the active pathways involved (Figure 6A). Our gene ontology (GO) molecular function analysis revealed translation dysregulation within cells upon knockdown of the lncRNA *TAAL*. Additionally, we observed the upregulation of proteins with ubiquitin ligase inhibitor and ubiquitin-protein transferase inhibitor activity (Figure 6B-C). To further validate our observation, we conducted a protein stability assay using cycloheximide (10μM) treatment under conditions of both knockdown and overexpression of lncRNA *TAAL*. Cycloheximide-mediated inhibition of translation resulted in decreased stability of the *Tie1* protein in both knockdown and overexpression scenarios. This indicates that the stability of *Tie1* is translation-dependent, with translation inhibition leading to reduced *Tie1* stability (Figure 6D).

**Figure 6:**
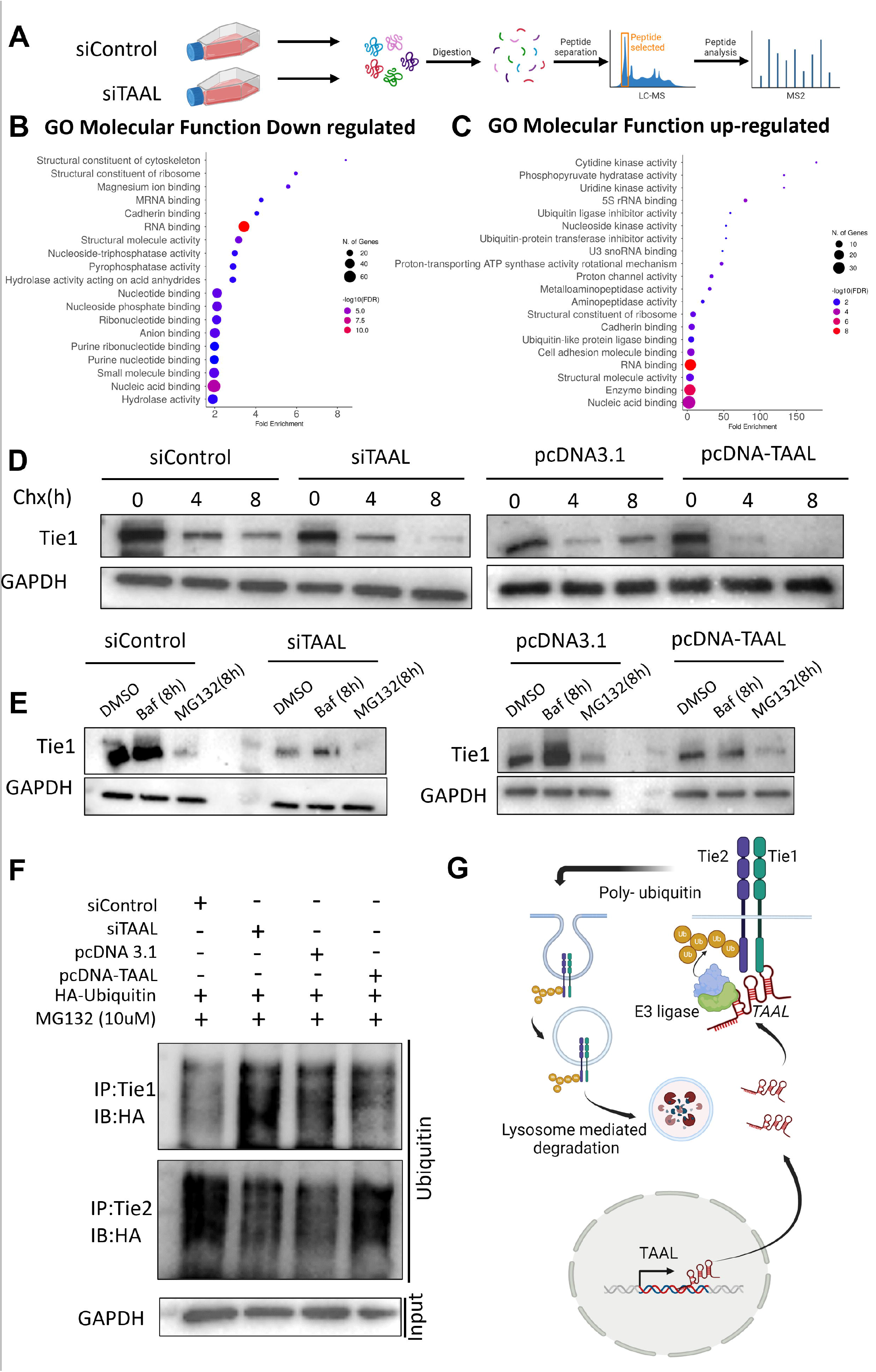
LncRNA *TAAL* controls the protein turnover of *Tie1* by modulating it ubiquitination. [A] Schematic of Protomics quntificatino pipeline for ceontrol and TAAL knockdown cells. [B] Dot plot representing Gene ontology molecular process for protein downregulated in *TAAL* knockdown cells upon comparing to the control. [C] Dot plot representing Gene ontology molecular process for protein up-regulated in *TAAL* knockdown cells upon comparing to the control. [D] Immunobloting of *Tie1* protein upon treatment of cells with translational inhibitor cycloheximide (Chx) for 0, 4, 8 hrs in control and *TAAL* knockdown or overexpressed cells. GAPDH was used as an input control. [E] Immunobloting of *Tie1* protein upon treatment of cells with degradation inhbitor MG132 and bafilomycin A1(Baf) for 8 hrs in control and *TAAL* knockdown or overexpressed cells. GAPDH was used as an input control. [F] Change in polyubiqutination of *Tie1* and *Tie2* protein in *TAAL* knockdown or overexpressed cells. Lysates of control and *TAAL* knockdown or overexpressed cells cotrnasfected with HA-Ubiquitin vector were immunoprecipitated with *Tie1* and *Tie2* antibody and immunoblotted with anti-HA antibody. [G] Hypothetical schematic illustrating the protective effect of lncRNA TAAL on Tie1 through polyubiquitination with heterotypic chains, which attract ubiquitin-like proteins and prevent lysosomal degradation.

Previous studies have suggested that *Tie* protein degradation occurs via the proteasomal pathway (Cheng & Hung, 2013; Wehrle *et al*, 2009) and as a receptor kinase, *Tie* may also be degraded through the lysosomal pathway(Gu *et al*, 2022; Zhao *et al*, 2022; Kreitman *et al*, 2018). To investigate this, we inhibited both the lysosomal and proteasomal pathways using the pharmacological drug bafilomycin A1 (50nM) for lysosome and MG132(10μM) for proteasome under conditions of knockdown and overexpression of lncRNA *TAAL*. Lysosomal pathway inhibition enhanced *Tie1* protein stability, whereas proteasomal pathway inhibition compromised stability and increased degradation (Figure 6E). These findings suggest that *Tie1* degradation involves both pathways, with proteasomal inhibition leading to increased lysosomal degradation.

Next, we examined the ubiquitination patterns of *Tie1* and *Tie2* proteins as its associated pathways were altered in proteomics studies. Knockdown of lncRNA *TAAL* resulted in increased ubiquitination of *Tie1* and decreased ubiquitination of *Tie2*. Conversely, overexpression of lncRNA *TAAL* led to decreased ubiquitination of *Tie1* and increased ubiquitination of *Tie2* (Figure 6F). These findings and insights from previous literature suggest that lncRNA *TAAL* may mediate the recruitment of specific ubiquitin ligases to *Tie1*, facilitating its ubiquitin-mediated lysosomal degradation. In the absence of lncRNA *TAAL*, the recruitment of appropriate ubiquitin ligases is impaired. Alternatively, we hypothesize that in the absence of *TAAL,* there could be the formation of heterotypic chains of ubiquitin on *Tie1* protein, leading to the binding of other ubiquitin-like proteins and thereby enhancing its stability (Figure 6G). Furthermore, the upregulation of ubiquitin ligase inhibitors under *TAAL* knockdown conditions significantly modulates the ubiquitin-degradation system, which is essential for the targeted degradation of *Tie1* proteins. The stability of both *Tie1* and *Tie2* proteins is enhanced by inhibiting ubiquitin ligases. These observations indicate that lncRNA *TAAL* plays a crucial role in regulating the ubiquitin-mediated lysosomal degradation of *Tie1/2*, thereby controlling protein turnover.

### lncRNA TAAL is negatively regulated in diabetic retinopathy

To investigate the disease association of lncRNA *TAAL*, we focused on retinopathy, a progressive neovascularization condition often resulting from diabetes. *Tie1/2* signaling pathway has been shown to play an important role in neovascularization during diabetic conditions. Hence we analyzed the expression of lncRNA *TAAL* in endothelial cells under hyperglycemic conditions.

High glucose treatments (25 mM and 50 mM) significantly downregulated lncRNA *TAAL* expression (Figure 7A). Data from the Noncode database revealed that lncRNA *TAAL* is present in extracellular vesicles secreted by HUVEC cells (Figure 7B) (Fang *et al*, 2018). Next, to validate our observation we examined the lncRNA *TAAL* expression in diabetic patients at various stages of retinopathy including patients having diabetes mellitus with no retinopathy (n=50), non-proliferative diabetic retinopathy (NPDR)(n=70), and proliferative diabetic retinopathy (PDR)(n=50) and compared it to control (n=60). We observed a significant enrichment of lncRNA *TAAL* in the blood samples of of NPDR (FC- 2.92; p-value < 0.001) and PDR (FC- 4.107; p-value < 0.001) patients compared to controls and other diabetic conditions (FC- 1.121; p-value - ns) (Figure 7C). Additionally, upon categorizing retinopathy based on its severity we observed that lncRNA TAAL was significantly higher in case of mild NPDR (FC 2.15, *p-*value <0.05), moderate NPDR (FC 3.99, *p-*value <0.001), severe NPDR (FC 2.66, *p-*value <0.05) and PDR (FC 4.1, *p-*value <0.00001) as compared to the healthy controls (Supply Fig 9A-B). Using normalized Ct values from NPDR patients versus controls, ROC curve analysis showed a significant correlation between NPDR and PDR with the lncRNA *TAAL* expression. We also computed the ROC area under the curve and observed a significant specificity and sensitivity for both NPDR (AUC- 0.6763; p-value - 0.004) and PDR (AUC- 0.7649; p-value < 0.0001) compared to control group (Figure 7 D-E). We next modeled the hyperglycemic conditions in HUVEC-hTert2 cells to mimic diabetic retinopathy and to understand the permeability defect associated with this condition. Hyperglycemic condition significantly increased FITC-dextran permeability across the endothelial monolayer (Figure 7F). We also observed decreased surface localization of VE-cadherin, a key adhesion molecule, under hyperglycemic conditions (Figure 7G). Next, we investigated the potential of lncRNA *TAAL* to modulate endothelial permeability. Overexpression of lncRNA *TAAL* significantly reduced permeability and restored VE-cadherin localization to the membrane under hyperglycemic conditions, as it decrease the *Tie1/2* levels inside the cells..

**Figure 7:**
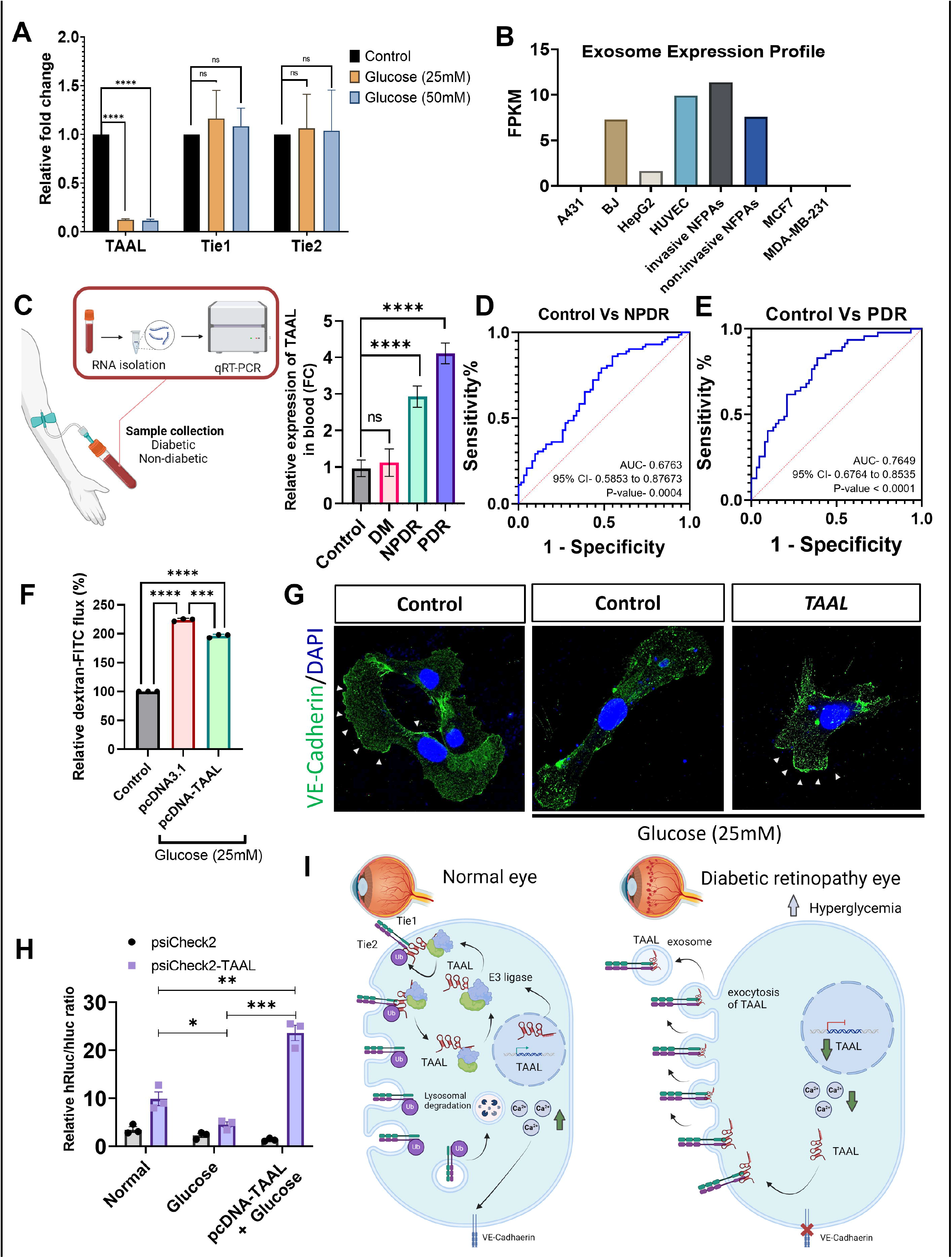
LncRNA *TAAL* plays a crucial role in regulating permeability and junctional dynamics in diabetic retinopathy. [A] Bar graph representing relative fold change levels of TAAL, Tie1, Tie2 upon treatment with hyperglysemia compared to control. ns- not significant; **** p-value ≤ 0.0001. (two-tailed unpaired t-test). [B] Expression profile of *TAAL* lncRNA in exosomes from different cell lines represented in as a bar graph. [C] Schematics of blood collection and quantification of lncRNA *TAAL* in diabetic and non-diabetics individuals. Bar graph represents relative fold change expression of lncRNA *TAAL* in blood samples of control, diabetic mellitus (DM), non-poliferative diabetic retinopathy (NPDR) and polyfrerative diabetic retinopathy (PDR) patients. [D-E] ROC curve analysis determines the sensitivity and specificity of lncRNA *TAAL* as diagnostic biomarker for NPDR or PDR patients with vascular dysfunction. [F] Complimentation of *TAAL* lncRNA in HUVEC hyperglysemic model reverts the increased permeability. Bar graph represents retalive dextran-FITC efflux in control untreated, control hyperglycemic and *TAAL* overexpressed in hyperglycemic cells. **** p-value ≤ 0.0001. (two-tailed unpaired t-test). [G] Complimentation of *TAAL* lncRNA in HUVEC hyperglysemic model reverts the localisation of VE-Cadhaerin at the junction of HUVEC cells. Confocal images of immunofluorescence VE-Cadhaerin in control and *TAAL* overexpressed cells under hyperglycemic condition (60X maginication). [H] Complimentation of *TAAL* lncRNA in HUVEC hyperglysemic model reverts the *in-vivo* expression of *TAAL* by increasing its promoter activity. Luciferase assay of lncRNA *TAAL* promoter in control untreated, control hypoglycemics and *TAAL* overexpressed hyperglycemic cells represented in a bar graph. * p-value ≤ 0.05; ** p-value ≤ 0.01; *** p-value ≤ 0.001 (two-tailed unpaired t-test). [I] Illustration showing lncRNA TAAL’s role in controlling vascular permeability and junctional dynamics through the regulation of Tie1 levels in normal cells. In hyperglycemic conditions, cells downregulate lncRNA TAAL expression and exocytose the residual TAAL transcripts into the bloodstream.

**Figure 8:**
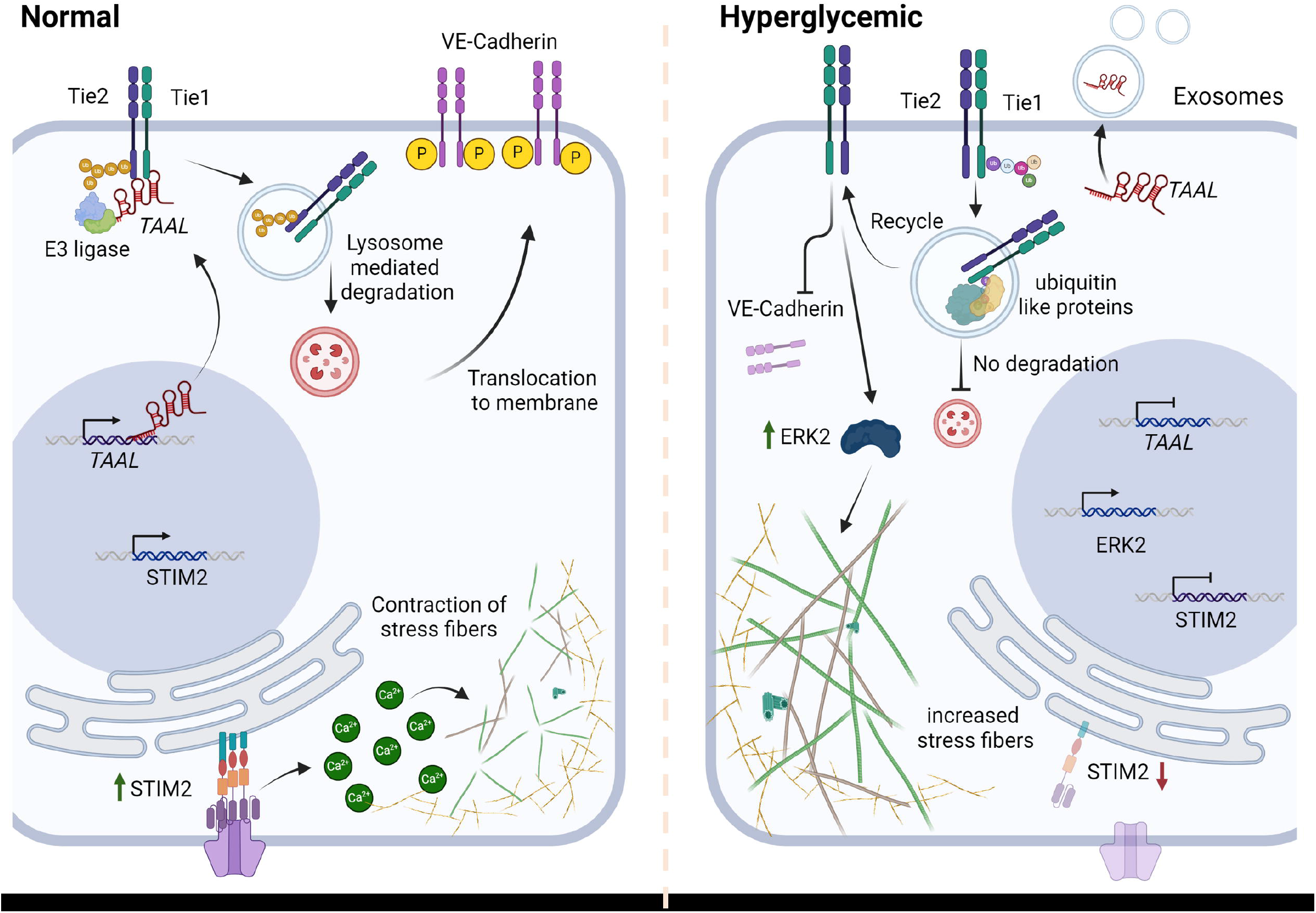
LncRNA *TAAL* controls the protein turnover of *Tie1* and modulates vascular permeability and angiogenic function.

To investigate the transcriptional regulation and impact of complementation on *TAAL* lncRNA under hyperglycemic conditions, we employed a luciferase reporter assay. This approach allowed us to determine whether the expression of lncRNA *TAAL* is regulated via a positive feedback loop upon complementation in hyperglycemia condition. We cloned the lncRNA *TAAL* promoter into the psiCHECK2 plasmid upstream of Renilla luciferase and performed assays in normal and hyperglycemic conditions. Hyperglycemia significantly decreased luciferase activity, suggesting high glucose inhibits lncRNA *TAAL* transcription. Furthermore, overexpression of lncRNA *TAAL* restored its transcription, indicating a positive feedback loop modulating its own expression. These data collectively underscore the significant role of lncRNA *TAAL* in modulating pathological conditions associated with diabetic retinopathy. We hypothesize that under hyperglycemic conditions, the transcription of lncRNA *TAAL* is downregulated by inhibiting its transcription. This process creates an environment that increases *Tie1* stability and promotes neovascularization, leading to pathological diabetic retinopathy. lncRNA *TAAL* has potential as a biomarker for predicting the progression from normal diabetes to diabetic retinopathy, as it is released into the bloodstream during retinopathy and can be detected by qRT-PCR. Furthermore, lncRNA *TAAL* exhibits high therapeutic potential for restoring normal conditions under hyperglycemia.

## Discussion

In this study, we utilized a protein-centric approach to identify interacting partners of the major endothelial receptor protein *Tie1* in HUVEC-hTert2 cells. We identified a novel, uncharacterized lncRNA named *TAAL*, whose modulation influences vascular function and integrity. Further investigations revealed that lncRNA *TAAL* modulates cellular calcium dynamics, affects cytoskeletal movements, and regulates VE-cadherin localization at cell junctions(Figure 3). Molecular analyses provided evidence that lncRNA *TAAL* controls the protein turnover of *Tie1* through the regulation of ubiquitin-mediated degradation and stability. Moreover, we identified a functional orthologue of human lncRNA *TAAL* in zebrafish, which does not exhibit any sequence conservation but has a common interacting partner *Tie1*. Knockdown of zebrafish *taal* resulted in angiogenic defects in the ISVs and SIVs (Figure 4 E-H). Rescue experiments demonstrated that human lncRNA *TAAL* can complement the function of zebrafish lncRNA *taal*, suggesting evolutionary conservation of function (Figure 4F). Our findings propose a negative regulatory role of *TAAL* over *Tie1,* thereby modulating its protein turnover. We further investigated the role of lncRNA *TAAL* in diabetic retinopathy a neovascular eye disease. We observed that the expression of lncRNA *TAAL* was significantly upregulated in patients with NPDR and PDR conditions (Figure 7C). Overexpression of lncRNA *TAAL* in a hyperglycemic model of HUVEC cells decreased permeability defects by restoring VE-cadherin at the membrane (Figure 7F-G). Additionally, overexpression of lncRNA *TAAL* induced a positive feedback loop, which enhanced the transcription of endogenous lncRNA *TAAL* (Figure 7H). Our investigation suggests that lncRNA *TAAL* is an important regulator of *Tie1* protein turnover and could be an effective therapeutic target for neovascular eye diseases.

Angiopoietin-Tie signaling is a crucial pathway for targeting various cancers and vascular eye diseases, specifically diabetic retinopathy (Saharinen *et al*, 2017; Joussen *et al*, 2021; Huang *et al*, 2010). *Tie1*, an orphan receptor, has been identified as an interacting receptor for angiopoietin-1 in a heterodimeric form with *Tie2*. In endothelial cells, the dynamics of *Tie1* and *Tie2* interaction and modulation play a significant role in their function. *Tie1* is a major protein associated with the angiogenic function of endothelial cells, particularly when *Tie2* levels are minimal. In the quiescent state, Tie2 levels are upregulated to maintain the survival of endothelial cells, while Tie1 levels remain low. During vascular remodeling, stalk cells exhibit a balance of Tie1 and Tie2 levels, underscoring the importance of both proteins (Savant *et al*, 2015). Modulating the interaction and levels of Tie1 and Tie2 *in vivo* holds promise for regulating endothelial cell functions. Inhibition of Angiopoietin-1 has been an effective strategy to modulate Tie1 levels and function during angiogenesis and disease conditions. Additionally, recent findings have identified *LECT2* as a receptor for *Tie1*, showing promising results in controlling angiogenesis and liver fibrosis in vivo (Xu *et al*, 2019). The development of second-generation targeted therapies for *Tie1* using antibodies has also demonstrated encouraging results in preclinical animal models, controlling angiogenesis and metastasis in cancer (Singhal *et al*, 2020; La Porta *et al*, 2018). With advancements in RNA modification and the emergence of RNA-based targeted therapies(Sparmann & Vogel, 2023; Saw & Song, 2024; Zhu *et al*, 2022), targeting a modulator of *Tie1* protein presents an intriguing approach. Our findings provide evidence that lncRNA *TAAL* can effectively modulate *Tie1* levels *in-vivo,* inhibiting downstream functions to control vascular remodeling. *TAAL* lncRNA serves as a promising candidate for RNA-based target therapy for modulating *Tie1* levels and its associated pathways in vascular diseases such as diabetic retinopathy and cancer metastasis.

LncRNAs have been shown to perform myriad functions within cells, primarily determined by their localization to the nucleus, chromatin, or cytoplasm. Cytoplasmic lncRNAs are of particular interest due to their accessibility for therapeutic targeting in various disease conditions. In the cytoplasm, lncRNAs are known to interact with proteins and regulate their functions and translation. Several studies, including those on lncRNAs such as *VEAL2, CamK-A, LINK-A, RP11-624L4.1, ECRAR, AGPG, and MetaLnc9 (LINC00963)*, have demonstrated that lncRNAs can interact with kinase proteins and modulate their signaling (Sehgal *et al*, 2021; Sang *et al*, 2018; Lin *et al*, 2016; Zhou *et al*, 2020; Chen *et al*, 2019; Liu *et al*, 2020; Yu *et al*, 2017). These interactions suggest roles for lncRNAs in acting as substrate-binding competitors, controlling localization, translation, and stability of kinase proteins. In our investigation, we found that the lncRNA *TAAL* controls the turnover of the receptor kinase *Tie1* protein. We hypothesize that in the absence of *TAAL* lncRNA, *Tie1* undergoes poly-ubiquitination with heterotypic chains of either K6 or K11 linkage, which are known to stabilize the protein (Tracz & Bialek, 2021; Hong *et al*, 2018; Abu Ahmad *et al*, 2021). These K6 and K11 chains of ubiquitin are known to evade lysosomal-mediated degradation by binding with ubiquitin-like proteins, thereby stabilizing the protein. Under normal conditions, poly-ubiquitination with either K63 or K48 linkage homotypic chains is associated with lysosomal degradation (Hong *et al*, 2018). The lncRNA *TAAL* may facilitate proper ubiquitination by recruiting E3 ligase to the *Tie1* protein. In hyperglycemic conditions, there is a complete shutdown of *TAAL* production in endothelial cells, and the remaining *TAAL* is removed via exocytosis. The absence of *TAAL* leads to poly-ubiquitination with heterotypic chains at the *Tie1* kinase domain, where ubiquitin-like proteins attach to these chains and protect them from degradation. Understanding the targets involved in proper ubiquitination of *Tie1* and the ubiquitin-like proteins that protect *Tie1* from degradation warrants further investigations. The protection of *Tie1 from degradation leads* to activation of *Tie1* downstream signaling cascades. This in turn results in change in cytoskeleton dynamics due to dysregulation of calcium dynamics, and alteration in localization of VE-cadherin on the surface of the cells (Figure 8). In this study we provide a detailed assessment and molecular function of lncRNA *TAAL* in vascular remodeling. Going forward, a comprehensive investigation is required to elucidate further the ubiquitination machinery involved in regulating *Tie1* expression.

Protein-centric targeted approaches in identifying lncRNA interactomes have shown great promise in selecting lncRNA modulators of disease-associated protein targets. These approaches are valuable not only for identifying orthologous lncRNAs across species that lack sequence conservation but also for understanding the multifaceted roles of lncRNAs in various biological processes(Ranjan *et al*, 2021; Diederichs, 2014; Ulitsky, 2016). Our recent work demonstrated that lncRNA *veal2* which was identified in zebrafish, has shown its interaction with *prkcb2*. A protein-centric pulldown technique was utilized to identify the human ortholog for zebrafish *veal2* without sequence conservation. *VEAL2* was shown to be a promising target and biomarker for proliferative diabetic retinopathy (PDR) patients, regulating vascular permeability (Sehgal *et al*, 2021). Similarly, lncRNA *JPX* a modulator of *XIST* lncRNA, which interacts with the *CTCF* protein, highlighted that this interaction is a conserved function of lncRNA *JPX* despite the absence of sequence conservation between human and mouse ortholog (Karner *et al*, 2020). In this study, we observed that despite the lack of sequence conservation between human and zebrafish lncRNA *TAAL*, *Tie1* is a conserved interacting partner, and *TAAL* functions similarly in both species. The conservation of lncRNA functions across species, even in the absence of sequence conservation, underscores the importance of considering alternative modes of conservation. Conservation based on motif or syntenic conservation has only identified a small proportion of orthologues and a large number of lncRNA remains unknown for their mode of conservation. Since many lncRNAs show little to no sequence conservation, focusing on protein-centric interactions can fill critical gaps in our understanding of lncRNA conservation and function. This targeted protein-centric approach can effectively identify functional conserved orthologous lncRNAs, providing insights into their evolutionary significance and biological roles.

In summary, we have identified a novel and uncharacterized lncRNA, *TAAL*, using a protein-centric pulldown approach. LncRNA *TAAL* plays a significant role in endothelial cell migration, permeability, and angiogenesis. By using *Tie1* as bait, we also identified the zebrafish ortholog of *TAAL*. Knockdown of zebrafish *taal* resulted in angiogenic defects in the intersegmental vessels (ISV) and subintestinal vessels (SIV), which were successfully rescued by human *TAAL*. Molecular investigations revealed that lncRNA *TAAL* regulates *Tie1* protein turnover by modulating its ubiquitination and subsequent degradation. Additionally, we observed that lncRNA *TAAL* levels were significantly upregulated in patients with diabetic retinopathy (NPDR & PDR), indicating its potential clinical relevance. Overexpression of lncRNA *TAAL* in hyperglycemic conditions restored endothelial cell permeability defects by enhancing the recruitment of VE-cadherin to membrane junctions. Our findings suggest that lncRNA *TAAL* plays a crucial role in regulating *Tie1* stability and function, highlighting its potential as a therapeutic target for vascular eye diseases.

## Structured Methods - Reagents and Tools Table

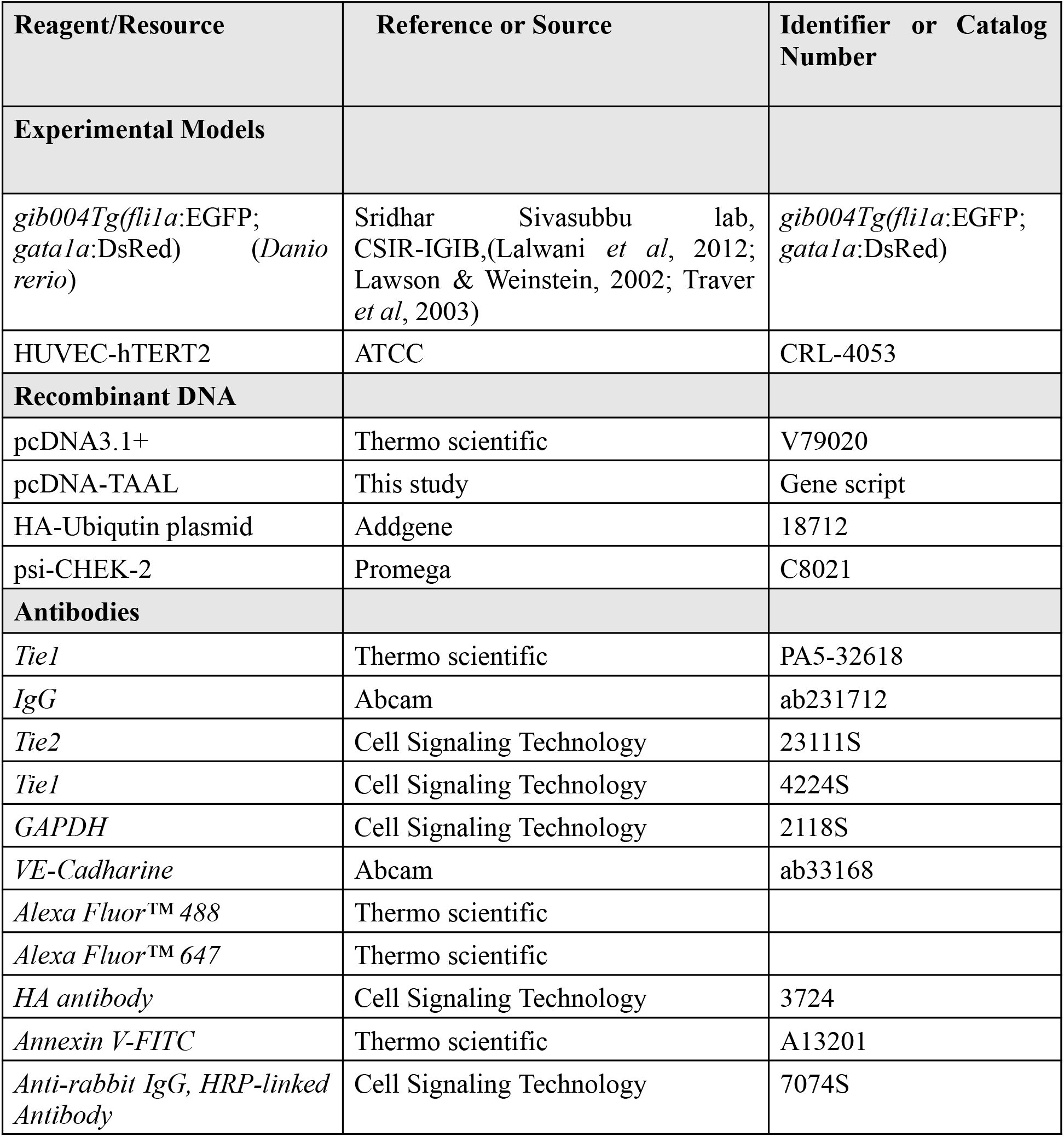

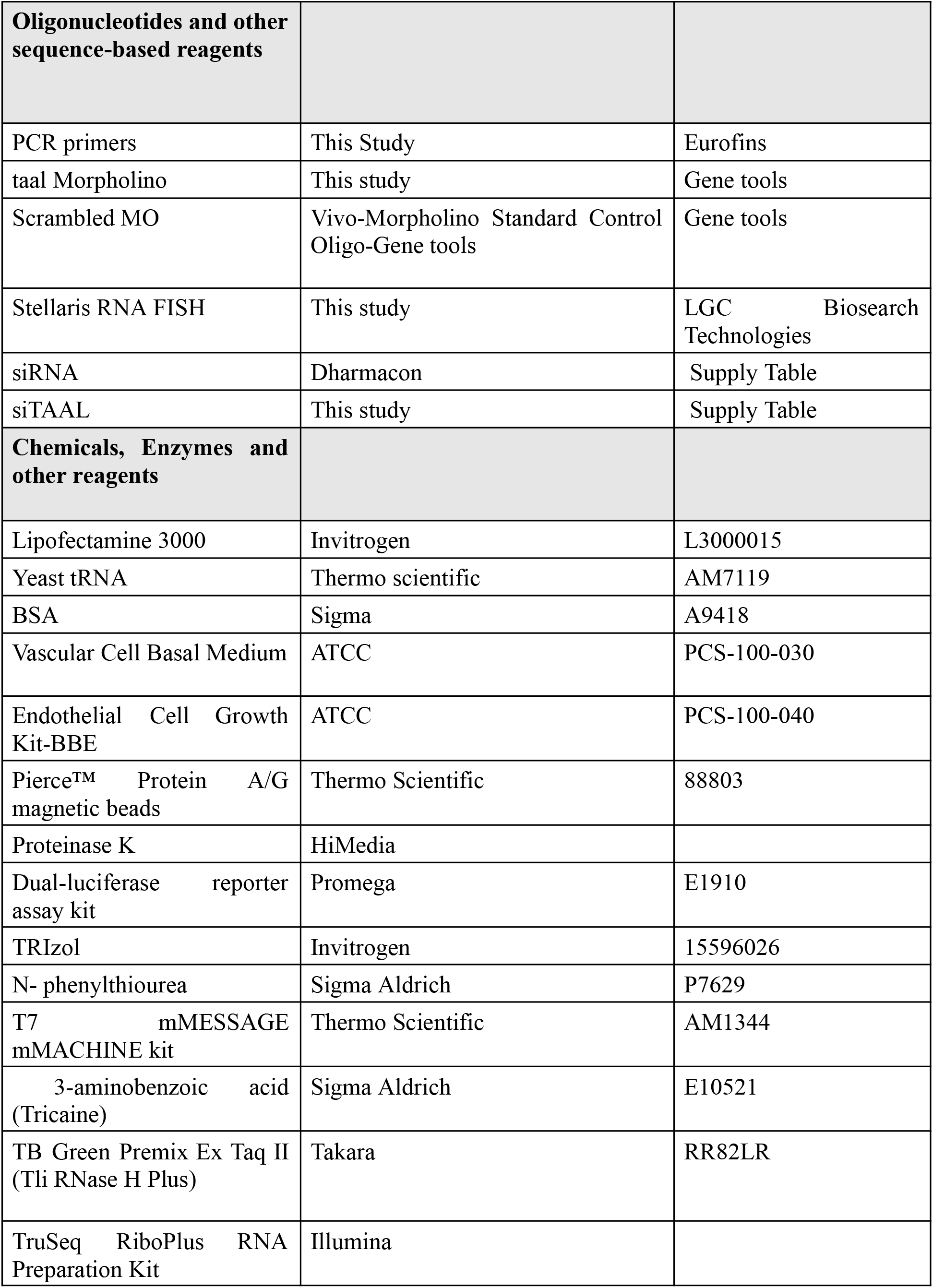

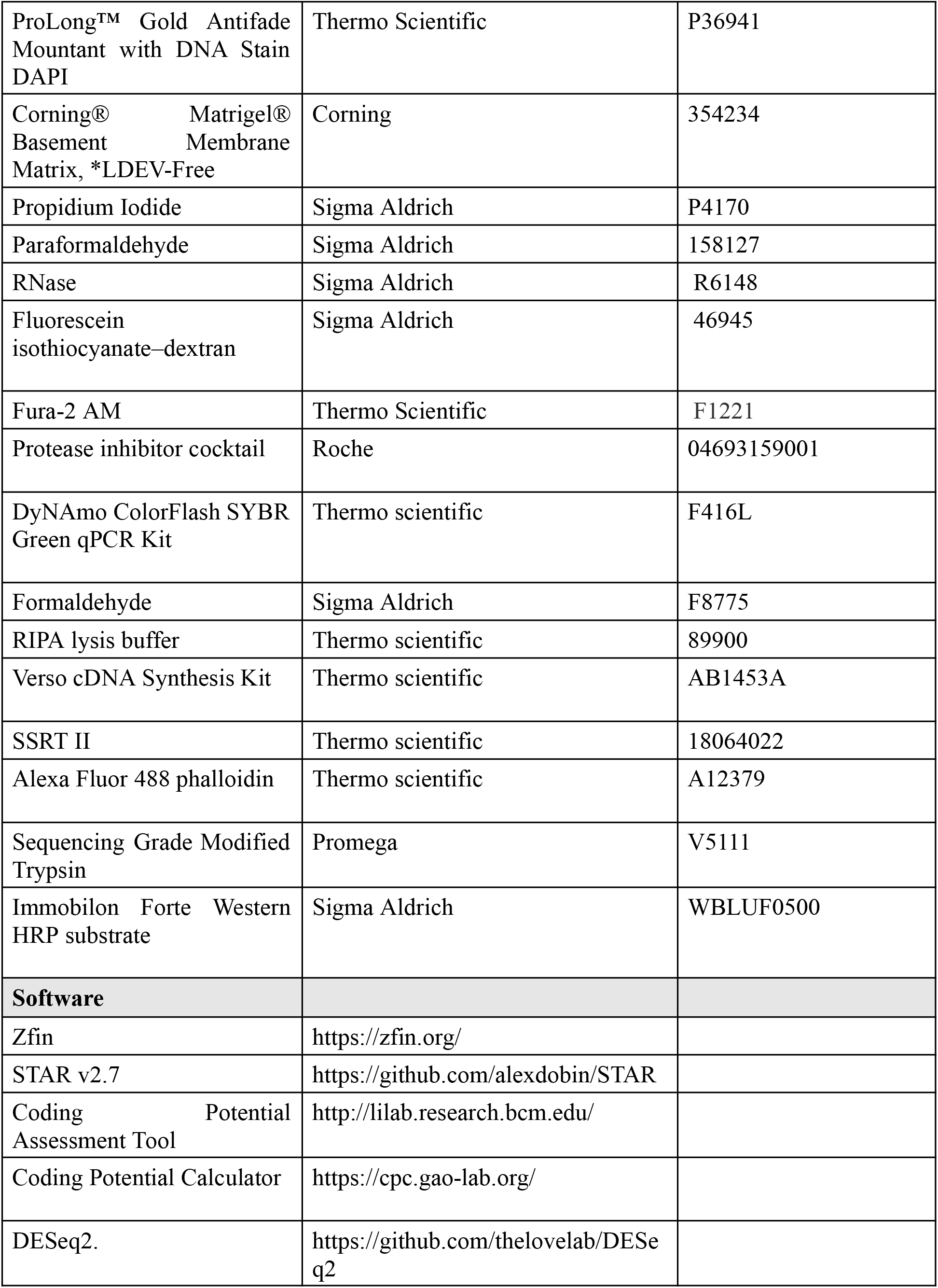

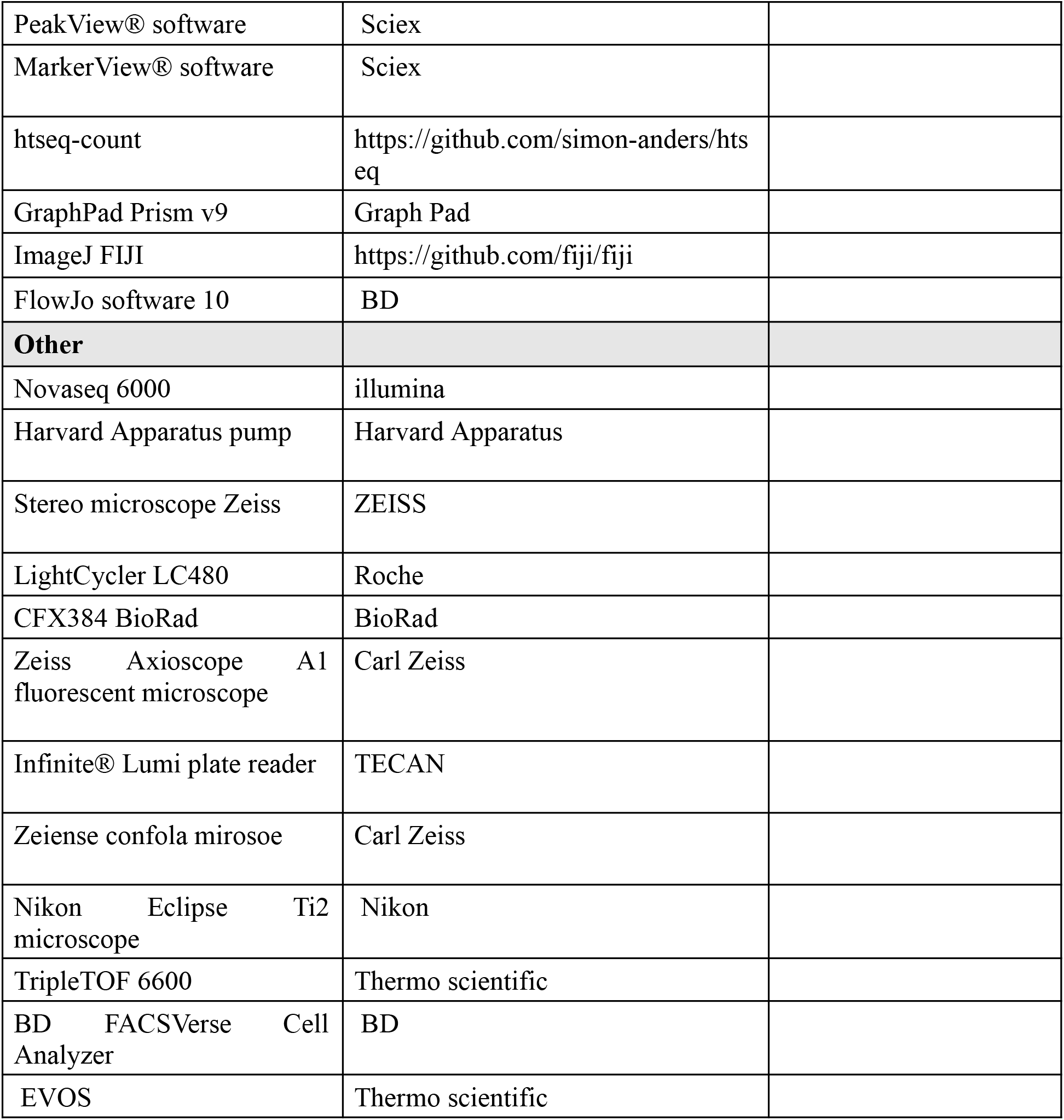

## Materials and Method

### Ethical approvals

Zebrafish used in this study is a double transgenic zebrafish (*Danio rerio*) *gib004Tg(fli1a*:EGFP; *gata1a*:DsRed) (Lalwani *et al*, 2012; Lawson & Weinstein, 2002; Traver *et al*, 2003). The zebrafish were cared for and maintained in-house following standard operating procedures approved by the Animal Ethics Committee of CSIR-Institute of Genomics and Integrative Biology, India (GENCODE-C-24). Special care was taken to ensure minimal distress to the animals.

The human study participants were recruited from Smt. Kannuri Santhamma Centre for vitreoretinal Diseases, L V Prasad Eye Institute, Hyderabad, India. The study was done according to the guidelines of Declaration of Helsinki and study was approved by Institutional Review Board (LEC-BHR-P-04-21-626).

### Cell line

We used Human umbilical vein endothelial cell lines HUVEC-hTert2 cell line (CRL-4053, ATCC) for our experiments (Karner *et al*, 2020; Takano *et al*, 2008). The cells were grown in Vascular Cell Basal Medium (PCS-100-030, ATCC) supplemented with Endothelial Cell Growth Kit-BBE (PCS-100-040, ATCC) incubated at 37 C with 5% CO2.

### RNA-Immunoprecipitation and RNA-sequencing

For RNA immunoprecipitation, we used ∼8 million HUVEC-hTert2 cells from two T75 flasks, and 500 zebrafish embryos at 24 hpf. UV cross-linking was performed according to a previously described protocol. Cell and zebrafish lysates were prepared using RIPA buffer supplemented with protease inhibitors. The cell lysate was incubated with protein A/G beads for pre-clearing for 2 hours at 4°C. Fresh protein A/G beads were blocked using 1x BSA and yeast tRNA for 1 hour at room temperature. The cell lysate, protein A/G beads, and the 1ug of primary antibody of *Tie1* (Thermo - PA5-32618) or *Tie2* (CST-23111S) were mixed together and incubated overnight on an orbital rotor at 4°C. The beads were then pulled down and washed with RIPA buffer. The beads were subsequently incubated in PNK buffer with proteinase K at 55°C for 45 minutes. Following incubation, the beads were dissolved in TRIzol, and RNA was isolated using the phenol-chloroform method. The isolated RNA was used for first-strand cDNA preparation using SSRT-II. The RNA-seq library was prepared using the TruSeq RiboPlus RNA Preparation Kit according to the manufacturer’s instructions. Finally, the library was sequenced on a NovaSeq 6000 platform.

The reads were aligned as previously described using STAR aligner to human reference genome hg38 and zebrafish reference genome Zv9 and reads were count using htseq -count function (Dobin *et al*, 2013; Anders *et al*, 2015; Ranjan *et al*, 2024).

### Western blot

Cells were lysed using NP40 lysis buffer supplemented with protease inhibitors and quantified using a BCA kit. The lysate was run on a 10% SDS-PAGE gel and transferred to a PVDF membrane. Primary antibodies for *Tie1* (CST-4224S) and *Tie2* (CST-23111S) were diluted to a concentration of 1:1000 and incubated overnight at 4°C. Secondary HRP-conjugated antibodies at 1:10000 dilution were incubated for 1 hour at room temperature. Immunoblotting was visualized using ECL under a chemiluminescence gel documentation system.

### Single molecular FISH and Immunofluorescent

We performed smFISH as per the manufacturer’s protocol (Biosearch Technologies). Both smFISH and co-immunofluorescence were conducted according to the provided instructions. Briefly, cells were grown on coverslips and fixed in 1% formaldehyde for 10 minutes, followed by three washes with 1X PBS. The cells were then permeabilized using 70% ethanol overnight at 4°C. After washing with 1X PBS three times, the cells were incubated in Buffer A for 10 minutes. The coverslips were then incubated with a hybridization buffer containing smFISH probes for *TAAL* lncRNA and primary antibody at a 1:200 dilution in a humidified chamber for 16 hours at 37°C. Following hybridization, the cells were washed with Wash Buffer A and incubated with a secondary antibody conjugated with Alexa Fluor for 30 minutes in the dark.

The cells were subsequently washed with Wash Buffer B, mounted on slides using Anti-fade Gold mounting media with DAPI, and fixed. Confocal imaging of the fixed slides was performed at 63X magnification with oil immersion. The data were processed and deconvolution was performed using FIJI ImageJ software and its plugins (Schindelin *et al*, 2012).

### Subcellular fraction

The subcellular fractionation of HUVEC-hTert2 was carried out following a modified version of the protocol described (Ten *et al*, 2012). In brief, T25 flask of HUVEC-hTert2 cells were incubated and homogenized in a hypotonic buffer consisting of 10 mM HEPES (pH 7.9), 1.5 mM MgCl₂, 10 mM KCl, 0.5 mM DTT, and a protease inhibitor. This mixture was then centrifuged at 1000 rpm for 5 minutes at 4°C. The supernatant, containing the cytoplasmic fraction, was separated, while the pellet, containing the nuclear fraction, was washed with the hypotonic buffer and collected. A small portion (1%) of each sample was set aside for western blot analysis, and the remainder was used for RNA extraction.

### Tube formation assay

Tube formation assay was performed on matrigel as previously described (Chatterjee *et al*, 2024). Briefly, the knockdown and overexpression of lncRNA TAAL in HUVEC-htert2 cells were taken for the assay. Matrigel was thawed and prepared as per the manufacturer’s protocol and was coated in a 48-well plate. 40,000 cells were taken for the assay from each condition and was loaded on top of the solidified matrigel with complete HUVEC mediate. The cells were incubated for 4-5 hrs at 37 ℃ and 5% CO2 and tube formation was imaged in EVOS or Nikkon microscope.

### Scratch wound assay

Scratch wound assay was performed as previously described (Sehgal *et al*, 2021). Briefly, HUVEC-hTert2 cells were transfected with siRNA and pcDNA3.1 plasmid targeting TAAL lncRNA for its knockdown and overexpression. Cells were grown in 24 well plates till the monolayer was formed, and before the experiment, the cells were incubated in HUVEC media without FBS. Using the p200 tip a scratch was made in the well, the media and cell debris were washed using 1 X PBS and incubated in HUVEC media with FBS for 24 hrs, and imaging was done in EVOS or Nikkon microscope.

### Annexin -V assay

Cell apoptosis was detected using an Annexin/PI-based assay. The HUVEC-htert2 cells were harvested after transient knockdown and overexpression of TAAL for 48 hours and resuspended in an annexin binding buffer. To this, Annexin V-FITC and Propidium Iodide (PI) were added in the dark for 10 min. Then, the data was acquired using BD FACSVerse Cell Analyzer (BD Bioscience) and the results were analyzed using FlowJo software 10.

### Cell cycle assay

To assess cell cycle progression upon knockdown and overexpression of *TAAL*, HUVEC-hTert2 cells were seeded in a 6-well plate. After 48 hours, cells were harvested and kept in 70% ethanol for 24 hours at -20 0 C. The next day, cells were treated with RNase at 37 0 C for 1 hour, then stained with Propidium Iodide (PI) in the dark for 30 minutes. The cell cycle distribution was detected using BD FACSVerse Cell Analyzer (BD Bioscience).

### Permeability assay

Permeability assay was performed as previously described (Sehgal *et al*, 2021). Briefly, lncRNA TAAL knockdown and overexpressed cells, along with control, were taken for the experiment. 1 million cells were seeded on the collagen-coated membrane’s upper chamber and incubated until the monolayer is formed. After the monolayer is formed, the media from the upper chamber is removed and supplemented with fresh media with fluorescein isothiocyanate (FITC) conjugated dextran (Sigma, USA) at 1mg/ml final concentration. The sample was incubated and after every 30 min, the media from the lower chamber was collected for 3 hours. UV spectrometer reading was taken on a monocular reader, and the FITC-dextrans efflux percentage was calculated.

### Ubiquitin assay

The ubiquitin pulldown was performed as previously described (Li *et al*, 2023). Briefly, HUVEC-hTert2 cells were cotransfected with siRNA / overexpression plasmid of TAAL lncRNA with HA-Ubiqutin plasmid (Addgene# 18712). The cells, after 48 hrs of treatment, were treated with MG132 for 8 hours and then harvested. The cells were washed with 1xPBS, and cell lysate was prepared using RIPA buffer (Thermo Scientific) having protease inhibitor and iodoacetamide. Cell lysate was precleared using protein A/G beads, and the primary antibody of target Tie1, and Tie2 was incubated with fresh protein A/G beads separately at room temperature for 2 hr. The cell lysate, proteinA/G bead conjugated primary antibody, was incubated overnight at 4 ℃ on the orbital rotor. ProteinA/G beads were pulled down and washed 3 times with RIPA buffer. The beads were dissolved in an SDS loading buffer and heated at 95 ℃ for 10 min. The sample was run on 10% SDS PAGE gel and was transferred on PVDF membrane. The blot was probed with HA antibody at 1:2000 dilution overnight at 4℃ and the secondary was incubated at room temperature for 1 hr. Immunoblotting was visualized using ECL under a chemiluminescence gel documentation system.

### Morpholino designing and zebrafish microinjection

Antisense morpholino oligonucleotide (MO) was designed to target the exons -intron boundaries of the lncRNA *taal*. The MO was ordered from GeneTools, redissolved in 300 ul to make 1mM stock, and stored in -20 ℃ for further use. We used 100uM concentration of each MO targeting both the introns and co-injected it into single-cell zebrafish or 2dpf zebrafish at SIV using FemtoJet injection system and Nikkon microscope. After injection, the embryos were transferred to (0.003%) PTU (N- phenylthiourea, Sigma Aldrich) water and further incubated at 28°C. Water was changed every 24 hrs, and dead embryos were discarded. Human lncRNA TAAL was amplified and converted into RNA using Invitro transcription kit T7mMassage mMachine (Thermo scientific). We injected 100ng/ul concentration into single-cell zebrafish or 2dpf zebrafish at SIV as described previously (Habeck *et al*, 2002; Sehgal *et al*, 2021).

### RT-qPCR

RNA isolation was done using TRIzol and phenol chloroform method and the samples were DNAses treated. RNA was converted to cDNA using an SSRTII (Thermo scientific) enzyme as instructed by the manufacturer. RT-qPCR was perfomred for taregt gene using TB Green SyBr (Takara) on Biorad CFX96 machine. The data was analyzed as described previously (Winer et al., 1999; Livak and Schmittgen, 2001).

### Proteomics

After knockdown using siRNA TAAL and siControl, cells were lysed using RIPA -lysis buffer. Proteins were precipitated with acetone chilled at -20°C. Supernatants were transferred, and four volumes of chilled acetone were added to it. After vortexing and incubation at -20°C for overnight, samples were centrifuged at 10,000 × g for 15 minutes at 4°C. The supernatant was discarded, the pellet air-dried, and resuspended in base buffer (Tris-Cl pH 8 (50 mM)). Protein concentration was determined by BCA assay. This sample was subjected to trypsin digestion (sequencing grade, Promega). 20ug of protein in 20 µl of base buffer was reduced with 2 µl of 25 mM DTT at 56°C for 30 minutes, followed by alkylation with 1 µl of 55 mM iodoacetamide at room temperature in the dark for 15 minutes. After adding 2 µl of trypsin (1 µg/µl), samples were incubated at 37°C for 18 hours. Digestion was terminated with 0.5% formic acid, and samples were vacuum evaporated at 30°C.

The peptides were fractioned into 8 fractions using a strong cation exchange column- ICAT (AB Sciex). Ellution of the peptide was done with a gradient concentration of ammonium formate buffer. The fractionated peptides were analyzed using a quadrupole-TOF hybrid mass spectrometer (TripleTOF 6600, AB Sciex, USA) coupled with a nano-LC system (Eksigent NanoLC-425) in SWATH mode for relative quantitation. Desalted peptides (5 µg) were initially loaded onto a trap-column (ChromXP C18CL 5 µm 120 Å, Eksigent) for additional desalting. Separation was performed on a reverse-phase C18 analytical column (ChromXP C18, 3 µm 120 Å, Eksigent) at a flow rate of 5 µl/minute with buffer A (water containing 0.1% formic acid) and buffer B (acetonitrile containing 0.1% formic acid). Each sample had a runtime of 50 minutes, followed by a 10-minute equilibration period before the next run. The raw data files generated by the instrument were imported into PeakView® software for further analysis. The prepared HUVEC-htert2 proteome ion library was utilized to identify MS/MS peaks, while relative quantitation of different proteins was conducted using MarkerView® software (AB Sciex).

### Luciferase assay

The promoter of lncRNA *TAAL* was cloned into the psiCHECK-2 plasmid using *NheI* and *KpnI* restriction enzymes, positioned in front of the renilla luciferase gene. HUVEC-hTert2 cells were transfected with siRNA and an overexpression plasmid for lncRNA *TAAL* along with psiCHEK-2 and psiCHEK-2-TAAL and grown under hyperglycemic (25 mM D-glucose) and normal conditions. Luciferase assays were performed after 48 hrs of transfection according to the manufacturer’s protocol (Promega).

### Calcium imaging

Calcium imaging was performed as previously described (Arora *et al*, 2021; Motiani *et al*, 2018). Briefly, HUVEC-hTert2 cells, after knockdown and overexpression of lncRNA *TAAL*, were plated on confocal dishes (SPL Life Sciences, Korea). After 48 hours of transfection, the cells were incubated with 4 μM fura-2 AM for 30 minutes at 37°C. The cells were then washed three times with HEPES-buffered saline solution (140 mM NaCl, 1.13 mM MgCl2, 4.7 mM KCl, 2 mM CaCl2, 10 mM D-glucose, and 10 mM HEPES; pH 7.4) and bathed for 5 minutes. For calcium imaging, a Nikon Eclipse Ti2 microscope coupled with a CoolLED pE-340 Fura light source and a high-speed PCO camera was used. Excitation was alternated at 340 and 380 nm, and emission was captured at 510 nm. The cells were treated with 10 μM thapsigargin to observe calcium release from the endoplasmic reticulum.

### Expression analysis in human blood samples

Blood samples were collected in EDTA-coated vials (for RNA isolation) from individuals with idiopathic macular holes and cataracts as normal controls for this study (n=60). Individuals with diabetic duration of more than 5 and no diabetic retinopathy served as diabetic controls for the study (n=50). Individuals with different forms of DR; mild NPDR (n=20), moderate NPDR (n=25), severe NPDR (n=25) and PDR (n=50) were recruited as cases for this study. Sample demographics are provided in the supply table 2. RNA was isolated from serum. The clot activator vials containing 1 ml blood samples were centrifuged at 1200 rpm for 10 minutes at 4 degrees Celsius and the supernatant was collected and stored at -80 degree Celsius until further use. The EDTA coated vials (containing 2 ml blood for RNA isolation) were subjected to series of incubations using the erythrocyte lysis buffer to get rid of red blood cell content and obtain a clear white pellet of white blood cells. RNA was extracted by the organic method using TRIzoL reagent from WBC pellet from blood. Quality and quantity were assessed by agarose gel and nanodrop spectrophotometer respectively. 500 ng of RNA was further reverse-transcribed to cDNA using the Verso cDNA Synthesis Kit (AB1453B). Semi-quantitative PCR was carried out using DyNAmo ColorFlash SYBR Green qPCR Kit (F-416L) and specific primers for lncRNA TAAL. β-actin was used as an endogenous control. Statistical analysis was performed using the student’s t-test.

## Data Availability

All the data has been provided in a supplementary file. Raw sequencing files will be shared upon request.

## Acknowledgments

This work was funded by DBT/Wellcome Trust India Alliance Fellowship (IA/I/19/2/504651) awarded to Rajender K Motiani. SA acknowledges Senior Research Fellowships from DBT. The authors want to acknowledge Dr. Soumya Sinha Roy and Dr. Barun Chatterjee for their valuable input in cell culture work. We also like to acknowledge Dr. Jyoti Tanwar for assistance and input with calcium imaging and Kriti Khare for assistance with confocal imaging. We also acknowledge the CSIR-IGIB common instrumentation facility, and the ATPC confocal facility, the FACS facility and all the LCSP lab members. Illustrations were made using BioRender.

## Author’s contribution

GR, SSB, RKM conceived and designed the experiment. GR, SA, LS, RB performed the experiments and bioinformatic analysis. SS, SC, IK performed clinical experiments. GR, SSB, and RKM analyzed the results. GR wrote the initial draft manuscript, and GR, SA, SSB, RKM edited the manuscript.

## References

1. Abu Ahmad Y, Oknin-Vaisman A, Bitman-Lotan E & Orian A (2021) From the Evasion of Degradation to Ubiquitin-Dependent Protein Stabilization. Cells 10

2. Anders S, Pyl PT & Huber W (2015) HTSeq--a Python framework to work with high-throughput sequencing data. Bioinformatics 31: 166–169

3. Arora S, Tanwar J, Sharma N, Saurav S & Motiani RK (2021) Orai3 Regulates Pancreatic Cancer Metastasis by Encoding a Functional Store Operated Calcium Entry Channel. Cancers 13

4. Augustin HG, Koh GY, Thurston G & Alitalo K (2009) Control of vascular morphogenesis and homeostasis through the angiopoietin-Tie system. Nat Rev Mol Cell Biol 10: 165–177

5. Bates DO & Curry FE (1997) Vascular endothelial growth factor increases microvascular permeability via a Ca(2+)-dependent pathway. American Journal of Physiology-Heart and Circulatory Physiology

6. Cao X, Li T, Xu B, Ding K, Li W, Shen B, Chu M, Zhu D, Rui L, Shang Z, et al (2023) Endothelial TIE1 Restricts Angiogenic Sprouting to Coordinate Vein Assembly in Synergy With Its Homologue TIE2. Arterioscler Thromb Vasc Biol 43: e323–e338

7. Carlantoni C, Allanki S, Kontarakis Z, Rossi A, Piesker J, Günther S & Stainier DYR (2021) Tie1 regulates zebrafish cardiac morphogenesis through Tolloid-like 1 expression. Dev Biol 469: 54–67

8. Chatterjee B, Fatima F, Seth S & Sinha Roy S (2024) Moderate Elevation of Homocysteine Induces Endothelial Dysfunction through Adaptive UPR Activation and Metabolic Rewiring. Cells 13

9. Cheng H-T & Hung W-C (2013) Inhibition of proliferation, sprouting, tube formation and Tie2 signaling of lymphatic endothelial cells by the histone deacetylase inhibitor SAHA. Oncol Rep 30: 961–967

10. Chen Y, Li X, Li B, Wang H, Li M, Huang S, Sun Y, Chen G, Si X, Huang C, et al (2019) Long Non-coding RNA ECRAR Triggers Post-natal Myocardial Regeneration by Activating ERK1/2 Signaling. Mol Ther 27: 29–45

11. Dalal PJ, Muller WA & Sullivan DP (2020) Endothelial Cell Calcium Signaling during Barrier Function and Inflammation. Am J Pathol 190: 535–542

12. D’Amico G, Korhonen EA, Waltari M, Saharinen P, Laakkonen P & Alitalo K (2010) Loss of Endothelial Tie1 Receptor Impairs Lymphatic Vessel Development-Brief Report. Arterioscler Thromb Vasc Biol

13. Diederichs S (2014) The four dimensions of noncoding RNA conservation. Trends Genet 30: 121–123

14. Dobin A, Davis CA, Schlesinger F, Drenkow J, Zaleski C, Jha S, Batut P, Chaisson M & Gingeras TR (2013) STAR: ultrafast universal RNA-seq aligner. Bioinformatics 29: 15–21

15. Duh EJ, Sun JK & Stitt AW (2017) Diabetic retinopathy: current understanding, mechanisms, and treatment strategies. JCI Insight 2

16. ENCODE Project Consortium, Moore JE, Purcaro MJ, Pratt HE, Epstein CB, Shoresh N, Adrian J, Kawli T, Davis CA, Dobin A, et al (2020) Expanded encyclopaedias of DNA elements in the human and mouse genomes. Nature 583: 699–710

17. Fang S, Zhang L, Guo J, Niu Y, Wu Y, Li H, Zhao L, Li X, Teng X, Sun X, et al (2018) NONCODEV5: a comprehensive annotation database for long non-coding RNAs. Nucleic Acids Res 46: D308–D314

18. Ferraro NM, Strober BJ, Einson J, Abell NS, Aguet F, Barbeira AN, Brandt M, Bucan M, Castel SE, Davis JR, et al (2020) Transcriptomic signatures across human tissues identify functional rare genetic variation. Science 369

19. Gavard J (2013) Endothelial permeability and VE-cadherin: a wacky comradeship. Cell Adh Migr 7: 455–461

20. Gu Y, Becker V, Qiu M, Tang T, Ampofo E, Menger MD & Laschke MW (2022) Brassinin Promotes the Degradation of Tie2 and FGFR1 in Endothelial Cells and Inhibits Triple-Negative Breast Cancer Angiogenesis. Cancers 14

21. Habeck H, Odenthal J, Walderich B, Maischein H, Schulte-Merker S &Tübingen 2000 screen consortium (2002) Analysis of a zebrafish VEGF receptor mutant reveals specific disruption of angiogenesis. Curr Biol 12: 1405–1412

22. Hamdollah Zadeh MA, Glass CA, Magnussen A, Hancox JC & Bates DO (2008) VEGF-mediated elevated intracellular calcium and angiogenesis in human microvascular endothelial cells in vitro are inhibited by dominant negative TRPC6. Microcirculation 15: 605–614

23. Hong S-Y, Kao Y-R, Lee T-C & Wu C-W (2018) Upregulation of E3 Ubiquitin Ligase CBLC Enhances EGFR Dysregulation and Signaling in Lung Adenocarcinoma. Cancer Res 78: 4984–4996

24. Huang H, Bhat A, Woodnutt G & Lappe R (2010) Targeting the ANGPT-TIE2 pathway in malignancy. Nat Rev Cancer 10: 575–585

25. Joussen AM, Ricci F, Paris LP, Korn C, Quezada-Ruiz C & Zarbin M (2021) Angiopoietin/Tie2 signalling and its role in retinal and choroidal vascular diseases: a review of preclinical data. Eye 35: 1305–1316

26. Kang Y-J, Yang D-C, Kong L, Hou M, Meng Y-Q, Wei L & Gao G (2017) CPC2: a fast and accurate coding potential calculator based on sequence intrinsic features. Nucleic Acids Res 45: W12–W16

27. Karner H, Webb C-H, Carmona S, Liu Y, Lin B, Erhard M, Chan D, Baldi P, Spitale RC & Sun S (2020) Functional Conservation of LncRNA JPX Despite Sequence and Structural Divergence. J Mol Biol 432: 283–300

28. Kreitman M, Noronha A & Yarden Y (2018) Irreversible modifications of receptor tyrosine kinases. FEBS Lett 592: 2199–2212

29. Lalwani MK, Sharma M, Singh AR, Chauhan RK, Patowary A, Singh N, Scaria V & Sivasubbu S (2012) Reverse genetics screen in zebrafish identifies a role of miR-142a-3p in vascular development and integrity. PLoS One 7: e52588

30. La Porta S, Roth L, Singhal M, Mogler C, Spegg C, Schieb B, Qu X, Adams RH, Baldwin HS, Savant S, et al (2018) Endothelial Tie1-mediated angiogenesis and vascular abnormalization promote tumor progression and metastasis. J Clin Invest 128: 834–845

31. Lawson ND & Weinstein BM (2002) In vivo imaging of embryonic vascular development using transgenic zebrafish. Dev Biol 248: 307–318

32. Lin A, Li C, Xing Z, Hu Q, Liang K, Han L, Wang C, Hawke DH, Wang S, Zhang Y, et al (2016) The LINK-A lncRNA activates normoxic HIF1α signalling in triple-negative breast cancer. Nat Cell Biol 18: 213–224

33. Liu J, Liu Z-X, Wu Q-N, Lu Y-X, Wong C-W, Miao L, Wang Y, Wang Z, Jin Y, He M-M, et al (2020) Long noncoding RNA AGPG regulates PFKFB3-mediated tumor glycolytic reprogramming. Nat Commun 11: 1507

34. Li X, Jiang W & Fang M (2023) An optimized protocol to detect ubiquitination modification of exogenous or endogenous proteins. STAR Protoc 4: 102650

35. Motiani RK, Tanwar J, Raja DA, Vashisht A, Khanna S, Sharma S, Srivastava S, Sivasubbu S, Natarajan VT & Gokhale RS (2018) STIM1 activation of adenylyl cyclase 6 connects Ca and cAMP signaling during melanogenesis. EMBO J 37

36. Mueller SB & Kontos CD (2016) Tie1: an orphan receptor provides context for angiopoietin-2/Tie2 signaling. J Clin Invest 126: 3188–3191

37. Pafumi I, Favia A, Gambara G, Papacci F, Ziparo E, Palombi F & Filippini A (2015) Regulation of Angiogenic Functions by Angiopoietins through Calcium-Dependent Signaling Pathways. Biomed Res Int 2015: 965271

38. Papapetropoulos A, Fulton D, Mahboubi K, Kalb RG, O’Connor DS, Li F, Altieri DC & Sessa WC (2000) Angiopoietin-1 inhibits endothelial cell apoptosis via the Akt/survivin pathway. J Biol Chem 275: 9102–9105

39. Pauli A, Valen E, Lin MF, Garber M, Vastenhouw NL, Levin JZ, Fan L, Sandelin A, Rinn JL, Regev A, et al (2012) Systematic identification of long noncoding RNAs expressed during zebrafish embryogenesis. Genome Res 22: 577–591

40. Phoenix KN, Yue Z, Yue L, Cronin CG, Liang BT, Hoeppner LH & Claffey KP (2022) PLCβ2 Promotes VEGF-Induced Vascular Permeability. Arterioscler Thromb Vasc Biol 42: 1229–1241

41. Ranjan G, Scaria V & Sivasubbu S (2024) Syntenic lncRNAs exhibit DNA regulatory functions with sequence evolution. bioRxiv

42. Ranjan G, Sehgal P, Sharma D, Scaria V & Sivasubbu S (2021) Functional long non-coding and circular RNAs in zebrafish. Brief Funct Genomics

43. Reinardy JL, Corey DM, Golzio C, Mueller SB, Katsanis N & Kontos CD (2015) Phosphorylation of Threonine 794 on Tie1 by Rac1/PAK1 Reveals a Novel Angiogenesis Regulatory Pathway. PLoS One 10: e0139614

44. Saharinen P, Eklund L & Alitalo K (2017) Therapeutic targeting of the angiopoietin-TIE pathway. Nat Rev Drug Discov 16: 635–661

45. Saharinen P, Eklund L, Miettinen J, Wirkkala R, Anisimov A, Winderlich M, Nottebaum A, Vestweber D, Deutsch U, Koh GY, et al (2008) Angiopoietins assemble distinct Tie2 signalling complexes in endothelial cell-cell and cell-matrix contacts. Nat Cell Biol 10: 527–537

46. Sahu SU, Visetsouk MR, Garde RJ, Hennes L, Kwas C & Gutzman JH (2017) Calcium signals drive cell shape changes during zebrafish midbrain-hindbrain boundary formation. Mol Biol Cell 28: 875–882

47. Sandoval R, Malik AB, Minshall RD, Kouklis P, Ellis CA & Tiruppathi C (2001) Ca(2+) signalling and PKCalpha activate increased endothelial permeability by disassembly of VE-cadherin junctions. J Physiol 533: 433–445

48. Sang L-J, Ju H-Q, Liu G-P, Tian T, Ma G-L, Lu Y-X, Liu Z-X, Pan R-L, Li R-H, Piao H-L, et al (2018) LncRNA CamK-A Regulates Ca-Signaling-Mediated Tumor Microenvironment Remodeling. Mol Cell 72: 601

49. Sato TN, Tozawa Y, Deutsch U, Wolburg-Buchholz K, Fujiwara Y, Gendron-Maguire M, Gridley T, Wolburg H, Risau W & Qin Y (1995) Distinct roles of the receptor tyrosine kinases Tie-1 and Tie-2 in blood vessel formation. Nature 376: 70–74

50. Savant S, La Porta S, Budnik A, Busch K, Hu J, Tisch N, Korn C, Valls AF, Benest AV, Terhardt D, et al (2015) The Orphan Receptor Tie1 Controls Angiogenesis and Vascular Remodeling by Differentially Regulating Tie2 in Tip and Stalk Cells. Cell Rep 12: 1761–1773

51. Saw PE & Song E (2024) Advancements in clinical RNA therapeutics: Present developments and prospective outlooks. Cell Rep Med 5: 101555

52. Schindelin J, Arganda-Carreras I, Frise E, Kaynig V, Longair M, Pietzsch T, Preibisch S, Rueden C, Saalfeld S, Schmid B, et al (2012) Fiji: an open-source platform for biological-image analysis. Nat Methods 9: 676–682

53. Sehgal P, Mathew S, Sivadas A, Ray A, Tanwar J, Vishwakarma S, Ranjan G, Shamsudheen KV, Bhoyar RC, Pateria A, et al (2021) LncRNA VEAL2 regulates PRKCB2 to modulate endothelial permeability in diabetic retinopathy. EMBO J 40: e107134

54. Shinde AV, Motiani RK, Zhang X, Abdullaev IF, Adam AP, González-Cobos JC, Zhang W, Matrougui K, Vincent PA & Trebak M (2013) STIM1 controls endothelial barrier function independently of Orai1 and Ca2+ entry. Sci Signal 6: ra18

55. Singhal M, Gengenbacher N, La Porta S, Gehrs S, Shi J, Kamiyama M, Bodenmiller DM, Fischl A, Schieb B, Besemfelder E, et al (2020) Preclinical validation of a novel metastasis-inhibiting Tie1 function-blocking antibody. EMBO Mol Med 12: e11164

56. Sparmann A & Vogel J (2023) RNA-based medicine: from molecular mechanisms to therapy. EMBO J 42: e114760

57. Stolz B & Bereiter-Hahn J (1988) Increase of cytosolic calcium results in formation of F-actin aggregates in endothelial cells. Cell Biol Int Rep 12: 321–329

58. Takano H, Murasawa S & Asahara T (2008) Functional and gene expression analysis of hTERT overexpressed endothelial cells. Biologics 2: 547–554

59. Ten S, Hodge K & I. A (2012) Dynamic proteomics: Methodologies and analysis. In Functional Genomics InTech

60. Tiruppathi C, Minshall RD, Paria BC, Vogel SM & Malik AB (2002) Role of Ca2+ signaling in the regulation of endothelial permeability. Vascul Pharmacol 39: 173–185

61. Tracz M & Bialek W (2021) Beyond K48 and K63: non-canonical protein ubiquitination. Cell Mol Biol Lett 26: 1

62. Traver D, Paw BH, Poss KD, Penberthy WT, Lin S & Zon LI (2003) Transplantation and in vivo imaging of multilineage engraftment in zebrafish bloodless mutants. Nat Immunol 4: 1238–1246

63. Tsai F-C & Meyer T (2012) Ca2+ pulses control local cycles of lamellipodia retraction and adhesion along the front of migrating cells. Curr Biol 22: 837–842

64. Tsai F-C, Seki A, Yang HW, Hayer A, Carrasco S, Malmersjö S & Meyer T (2014) A polarized Ca2+, diacylglycerol and STIM1 signalling system regulates directed cell migration. Nat Cell Biol 16: 133–144

65. Ulitsky I (2016) Evolution to the rescue: using comparative genomics to understand long non-coding RNAs. Nat Rev Genet 17: 601–614

66. Wang L, Park HJ, Dasari S, Wang S, Kocher J-P & Li W (2013) CPAT: Coding-Potential Assessment Tool using an alignment-free logistic regression model. Nucleic Acids Res 41: e74

67. Wehrle C, Van Slyke P & Dumont DJ (2009) Angiopoietin-1-induced ubiquitylation of Tie2 by c-Cbl is required for internalization and degradation. Biochem J 423: 375–380

68. Wong TY, Cheung CMG, Larsen M, Sharma S & Simó R (2016) Diabetic retinopathy. Nature Reviews Disease Primers 2: 1–17

69. Xu M, Xu H-H, Lin Y, Sun X, Wang L-J, Fang Z-P, Su X-H, Liang X-J, Hu Y, Liu Z-M, et al (2019) LECT2, a Ligand for Tie1, Plays a Crucial Role in Liver Fibrogenesis. Cell 178: 1478–1492.e20

70. Yang H, Luan Y, Liu T, Lee HJ, Fang L, Wang Y, Wang X, Zhang B, Jin Q, Ang KC, et al (2020) A map of cis-regulatory elements and 3D genome structures in zebrafish. Nature 588: 337–343

71. Yu T, Zhao Y, Hu Z, Li J, Chu D, Zhang J, Li Z, Chen B, Zhang X, Pan H, et al (2017) MetaLnc9 Facilitates Lung Cancer Metastasis via a PGK1-Activated AKT/mTOR Pathway. Cancer Res 77: 5782–5794

72. Zhao L, Zhao J, Zhong K, Tong A & Jia D (2022) Targeted protein degradation: mechanisms, strategies and application. Signal Transduct Target Ther 7: 113

73. Zhou L, Liu R, Liang X, Zhang S, Bi W, Yang M, He Y, Jin J, Li S, Yang X, et al (2020) lncRNA RP11-624L4.1 Is Associated with Unfavorable Prognosis and Promotes Proliferation via the CDK4/6-Cyclin D1-Rb-E2F1 Pathway in NPC. Mol Ther Nucleic Acids 22: 1025–1039

74. Zhu Y, Zhu L, Wang X & Jin H (2022) RNA-based therapeutics: an overview and prospectus. Cell Death Dis 13: 644

