## Supplementary material for "LncRNA *TAAL* is a Modulator of *Tie1*-Mediated Vascular Function in Diabetic Retinopathy": Supply

### Supplementary Data

**Table 1**: Significant RNA seq reads human and zebrafish
The Excel file will be provided upon request

**Table 2:** Demographics for the samples used for expression analysis of lncRNA TAAL in the blood samples; &= DM duration non-significant with the diabetic group. # age difference non-significant with the controls.

| | **Control** | **DM** | **Mild NPDR** | **Moderate NPDR** | **Severe NPDR** | **PDR** |
|---|---|---|---|---|---|---|
| **Age** | 64±7 | 59±7.8# | 60.65±7.9# | 60.5±7# | 58.5±4# | 53.66±6.8# |
| **Gender (F/M)** | 27/32 | 17/31 | 05/17 | 05/21 | 07/18 | 11/38 |
| **DM duration** | | 12.35±7.37 | 12.1±5.4& | 14.1±6.19& | 12.3±4.5& | 12.8±6.7& |

**Table 3**: Raw reads patient data
The Excel file will be provided upon request

**Table 4** : Primers used in the study

| **Name** | **Seq 5'-3'** | **Length** |
|---|---|---|
| TAAL Full length F | AGGAAGAGGAGGATTCCGAT | 20 |
| TAAL Full length R | AATTGTTCATGGGAGATCAT | 20 |
| T7+TAAL Full length F | TAATACGACTCACTATAGGGAGGAAGAGGAGGATTCCGAT | 40 |
| TAAL rtpcr F | GATGACAATGATGGCCTGTG | 20 |
| TAAL rtpcr R | AGAAAGTAGCTGCTGCGTCT | 20 |
| VE-Cad F | GTGTTCACGCATCGGTTGTT | 20 |
| VE-Cad R | CCCCTTCAGGATTTGGTACA | 20 |
| VEGFR2 F | ATGCTGGCATGGTCTTCTGT | 20 |
| VEGFR2 R | ATGAGACGGACTCAGAACCA | 20 |
| MAPK1 F (ERK1) | TTCCAACCTGCTGCTCAACA | 20 |
| MAPK1 R (ERK1) | GAACCCTGTGTGATCATGGT | 20 |
| MAPK3 F (ERK2) | GATCTAAAGCCCTCCAACCT | 20 |
| MAPK3 R (ERK2) | GTCGATGGACTTGGTATAGC | 20 |
| RAC1 F | ACCGGTGAATCTGGGCTTAT | 20 |
| RAC1 R | ATGCAGGACTCACAAGGGAA | 20 |

| | | |
|---|---|---|
| STIM1 F | CCAGAGCCTCAGCCATAGTC | 20 |
| STIM1 R | GGTCTTCCCTCAGGAACTCATC | 22 |
| STIM2 F | CCCTCACCACCCGCAACA | 0 |
| STIM2 R | GATGTGTGGCGAGGTTAAGGC | 0 |
| ORAI1 F | AGGTGATGAGCCTCAACGAG | 20 |
| ORAI1 R | CTGATCATGAGCGCAAACAG | 20 |
| ORAI2 F | GCAGCTACCTGGAACTGGTC | 20 |
| ORAI2 R | CGGGTACTGGTACTGCGTCT | 20 |
| ORAI3 F | GAGTGACCACGAGTACCCACC | 21 |
| ORAI3 R | GGGTACCATGATGGCTGTGG | 20 |
| Zf-taal Morholino 1 | ATCAGTTCTACCTGCTTCCTGA | 22 |
| Zf-taal Morholino 2 | ATCAGCTTTACCTGCTTCCTGAGC | 24 |
| GAPDH F | AACTGCTTAGCACCCCTGGC | 20 |
| GAPDH R | ATGACCTTGCCCACAGCCTT | 20 |
| siRNA-TAAL | UCAUUAGAGAGCCCAAAUCUU | 21 |

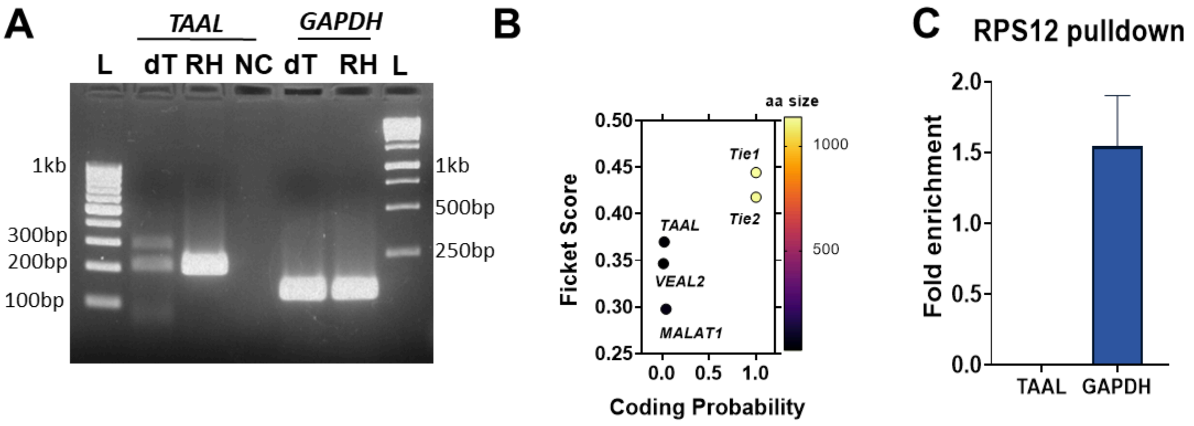


**Supplementary Figure 1:**

[A] Image showing agarose gel electrophoresis of the PCR product derived from *TAAL* lncRNA and *GAPDH*, amplified from cDNA synthesized using both oligo dT(dT) and random hexamer primers (RH).

[B]Coding potentiality scores were calculated using CPC2 for *TAAL* and other lncRNA and protein-coding genes (*Tie1 and Tie2*).

[C] Bar plot representing fold enrichment of *TAAL and GAPDH* upon RNA-immunoprecipitation of Ribosomal 40S subunit (RPS12) by qPCR; Data from 3 independent biological replicates plotted as mean fold change ± SEM.

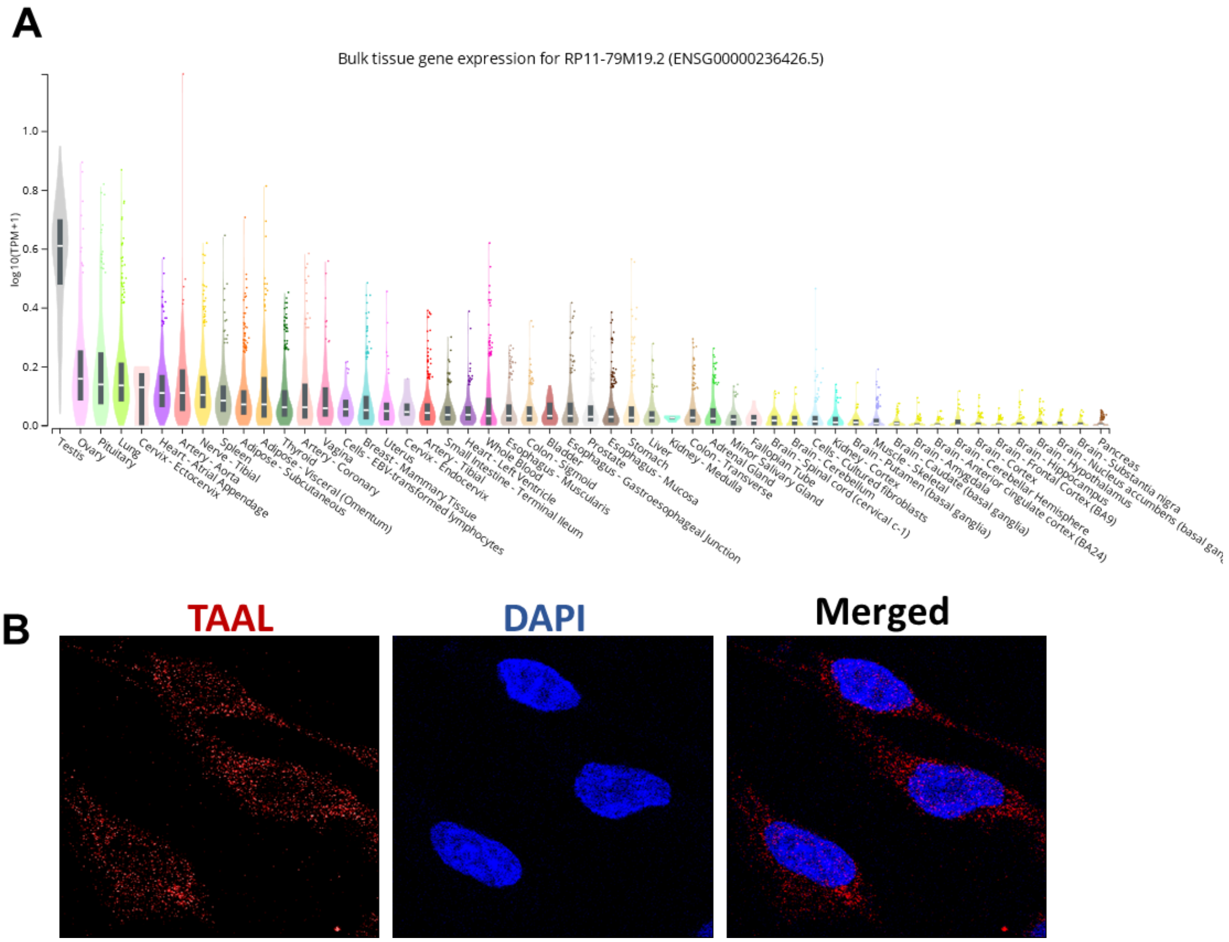


**Supplementary Figure 3:**

[A] Expression profile of *TAAL* lncRNA across different tissues in GTEx v8 database.

[B] Confocal image representing single molecular *FISH* (*smFISH*) of *TAAL* lncRNA. At 60x magnification.

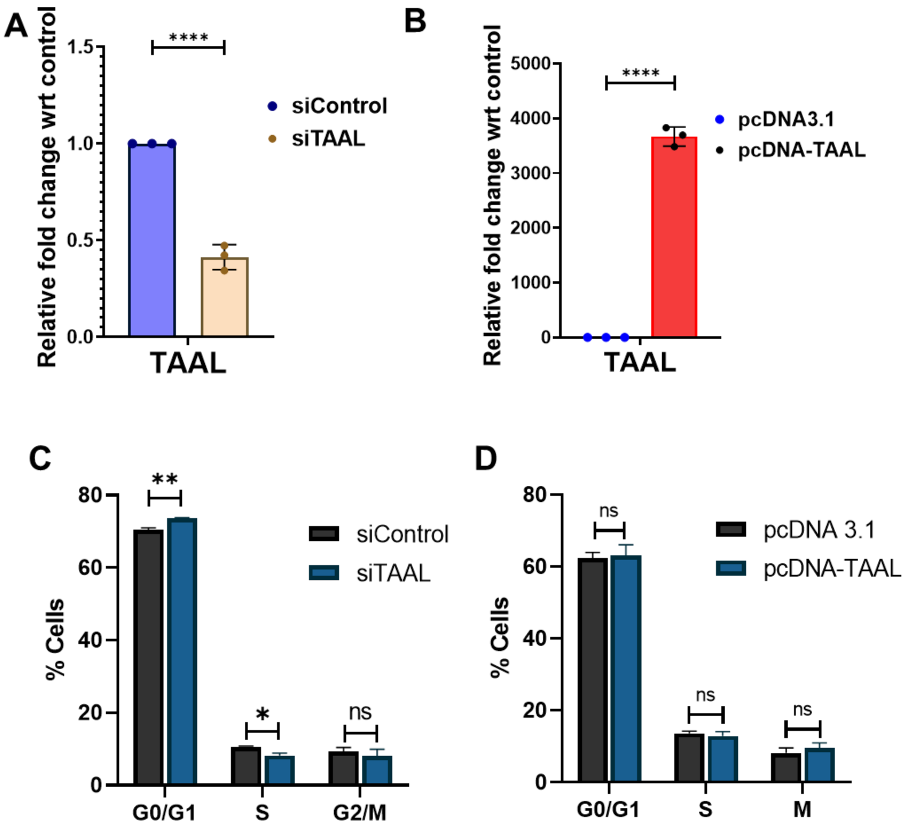


**Supplementary Figure 3:**

[A] Bar plot representing fold enrichment of lncRNA *TAAL* upon knockdown with siControl and siTAAL after 48hrs of transfection. Data from 3 independent biological replicates plotted as mean fold change ± standard deviation. **** p-value ≤ 0.0001. (two-tailed unpaired t-test)

[B] Bar plot representing fold enrichment of lncRNA *TAAL* upon overexpression with pcDNA3.1 and pdDNA-TAAL plasmid after 48hrs of transfection. Data from 3 independent biological replicates plotted as mean fold change ± standard deviation. **** p-value ≤ 0.0001. (two-tailed unpaired t-test)

[C] Bar plot illustrating the cell cycle distribution of cells 48 hours post-transfection with siControl and siTAAL. Cells were stained with propidium iodide (PI) and analyzed by FACS to determine cell cycle phases. Data from 3 independent biological replicates

[D]Bar plot illustrating the cell cycle distribution of cells 48 hours post-transfection with pcDNA3.1 and pdDNA-TAAL plasmid. Cells were stained with propidium iodide (PI) and analyzed by FACS to determine cell cycle phases. Data from 3 independent biological replicates

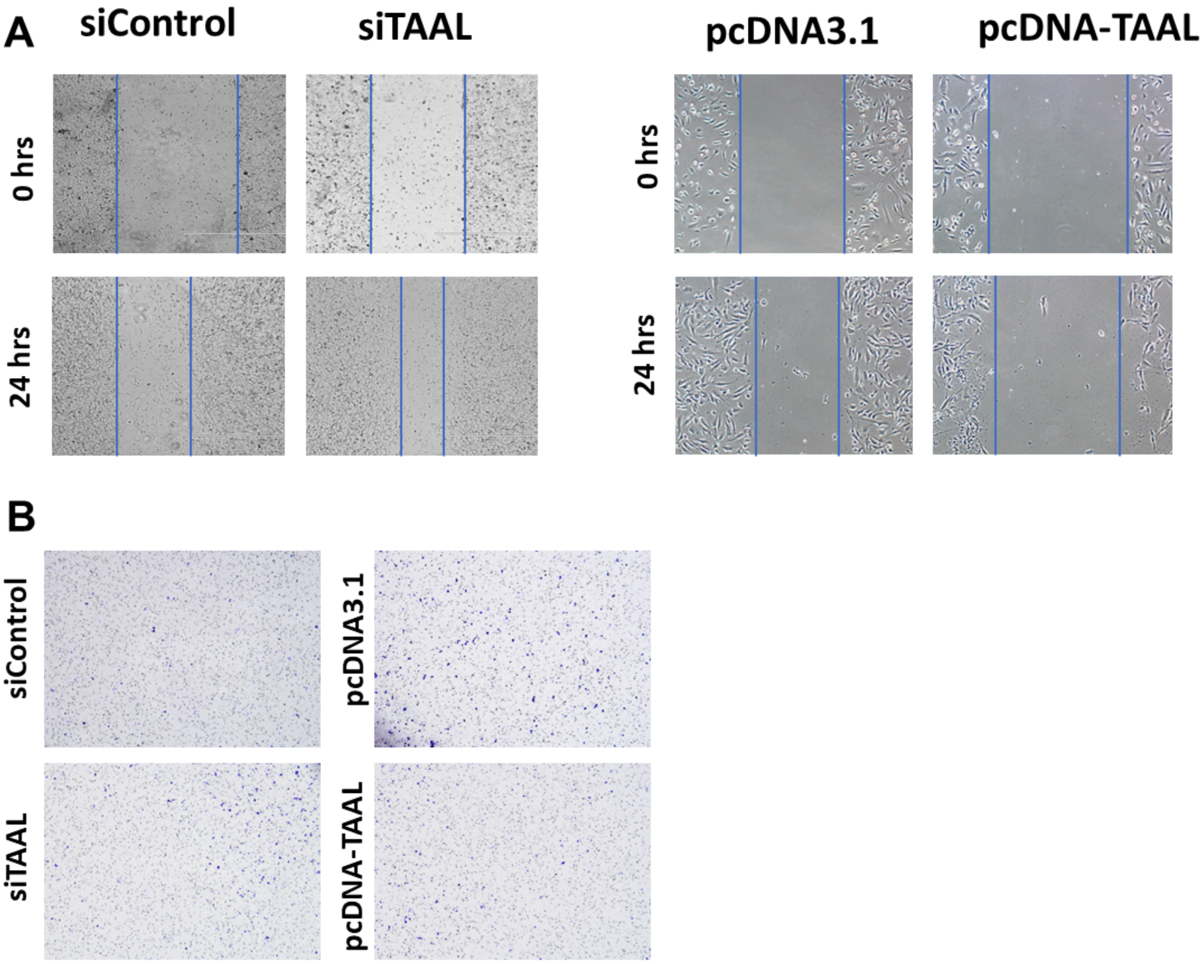


**Supplementary Figure 4: [A]** Raw image for wound closer rate in scratch wound assay after 24 hrs in control and *TAAL* knockdown or overexpressed cells.

[B] Raw image of cells migration in the transwell assay compared in control and *TAAL* knockdown or overexpressed cells.

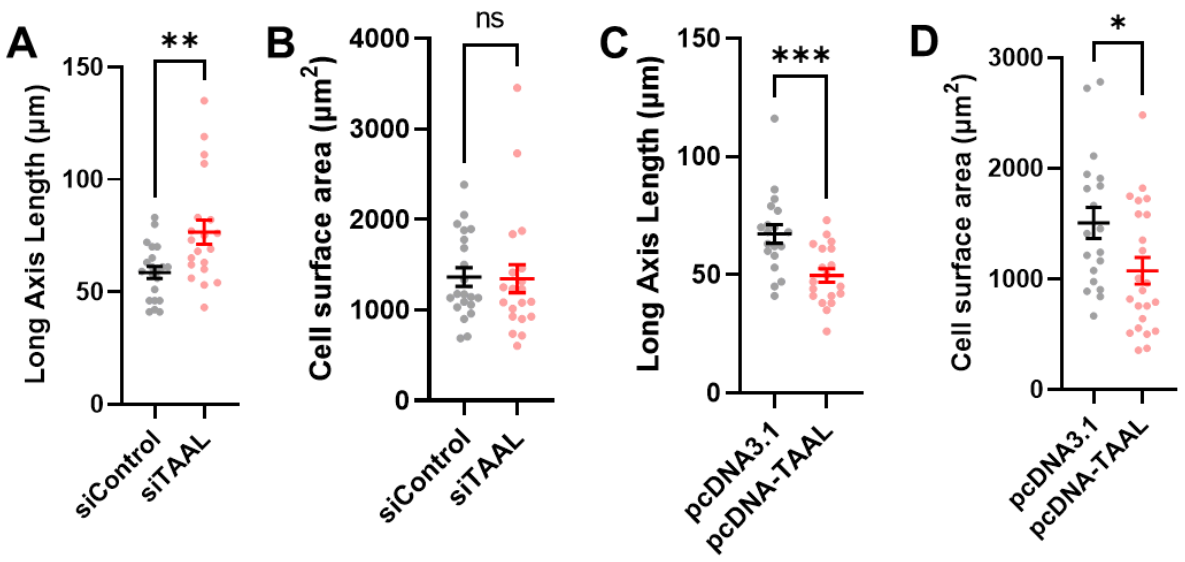


**Supplementary Figure 5:**
[A] Dot plot representing quantification of long axis in control and *TAAL* knockdown cells. Each dot represents an individual cell. ** p-value ≤ 0.01 (two-tailed unpaired t-test).
[B] Dot plot representing quantification of cell surface area (in $\mu m^2$) in control and *TAAL* knockdown cells. Each dot represents an individual cell ns-not significant (two-tailed unpaired t-test).
[C] Dot plot representing quantification of long axis in control and *TAAL* knockdown cells. Each dot represents an individual cell. *** p-value ≤ 0.001 (two-tailed unpaired t-test).
[A] Dot plot representing quantification of cell surface area (in $\mu m^2$) in control and *TAAL* knockdown cells. Each dot represents individual cell * p-value ≤ 0.05 (two-tailed unpaired t-test).

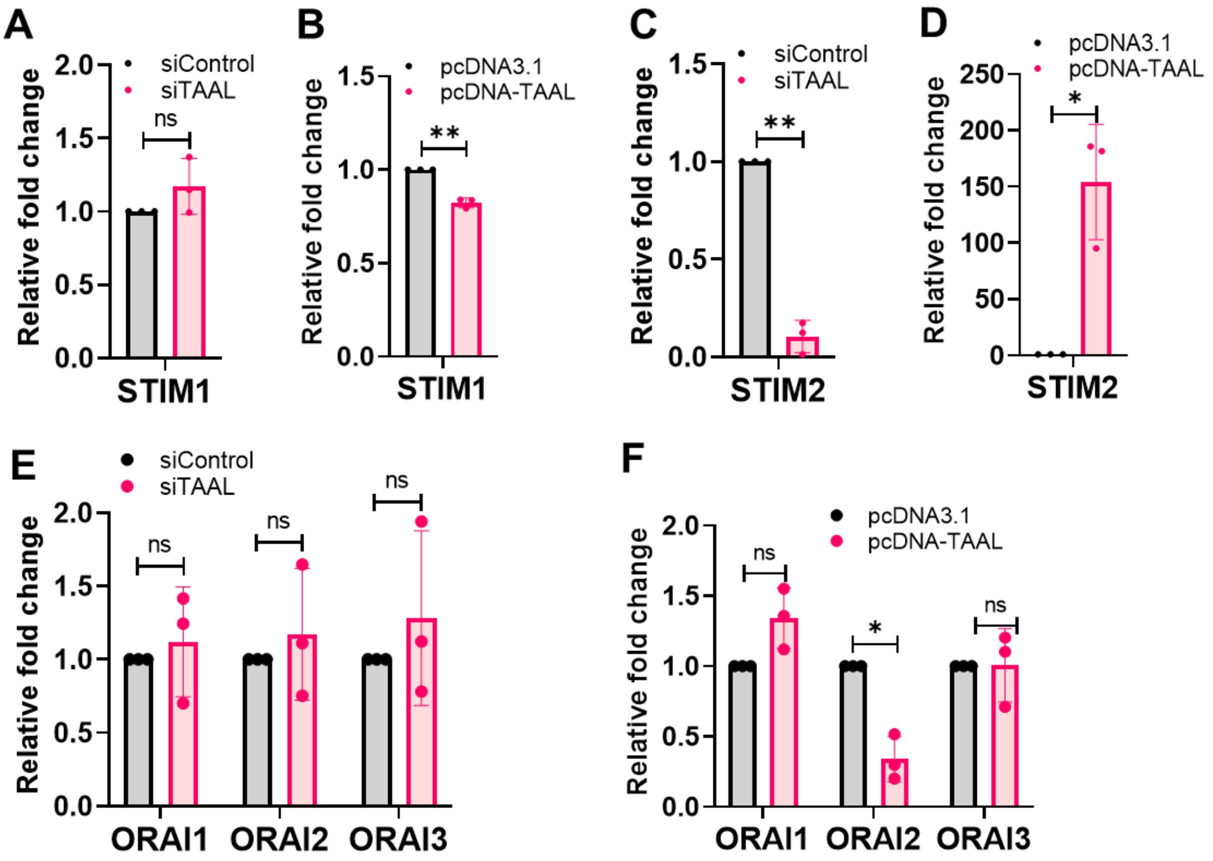


**Supplementary Figure 6:**

[A, C, E] Bar plot representing fold change of STIM1, STIM2, ORAI1, ORAI2 and ORAI3 upon knockdown with siControl and siTAAL after 48hrs of transfection. Data from 3 independent biological replicates plotted as mean fold change ± standard deviation. ns- not significant; ** p-value ≤ 0.01. (two-tailed unpaired t-test)

[B, D, F] Bar plot representing fold change of STIM1, STIM2, ORAI1, ORAI2 and ORAI3 upon overexpression with pcDNA3.1 and pcDNA-TAAL after 48hrs of transfection. Data from 3 independent biological replicates plotted as mean fold change ± standard deviation. ns- not significant; * p-value ≤ 0.05; ** p-value ≤ 0.01. (two-tailed unpaired t-test)

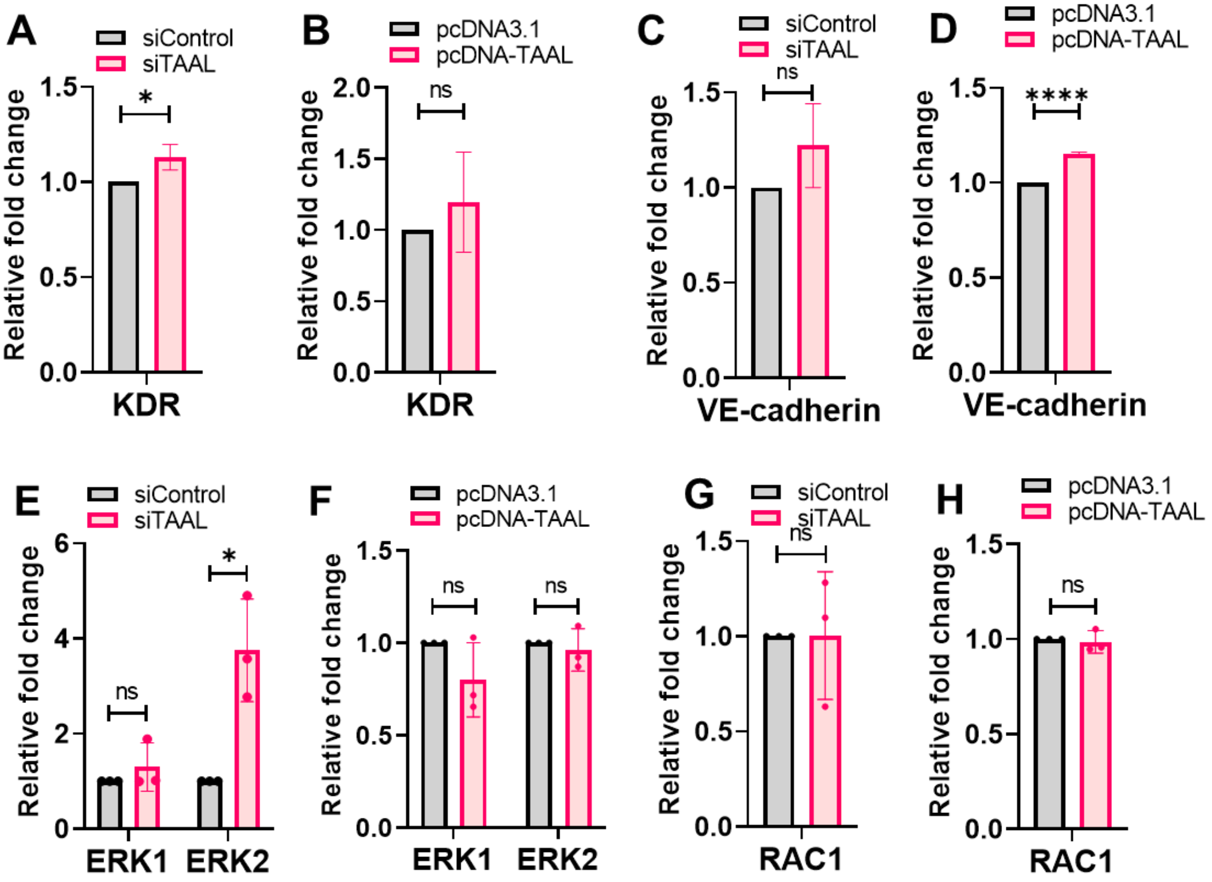


**Supplementary Figure 7:**

[A, C, E, G] Bar plot representing fold change of KDR, VE-Cadharin, ERK1, ERK2 and RAC1 upon knockdown with siControl and siTAAL after 48hrs of transfection. Data from 3 independent biological replicates plotted as mean fold change ± standard deviation. ns- not significant; * p-value ≤ 0.05. (two-tailed unpaired t-test)

[B, D, F, H] Bar plot representing fold change of KDR, VE-Cadharin, ERK1, ERK2 and RAC1 upon overexpression with pcDNA3.1 and pcDNA-TAAL after 48hrs of transfection. Data from 3 independent biological replicates plotted as mean fold change ± standard deviation. ns- not significant; **** p-value ≤ 0.0001. (two-tailed unpaired t-test)

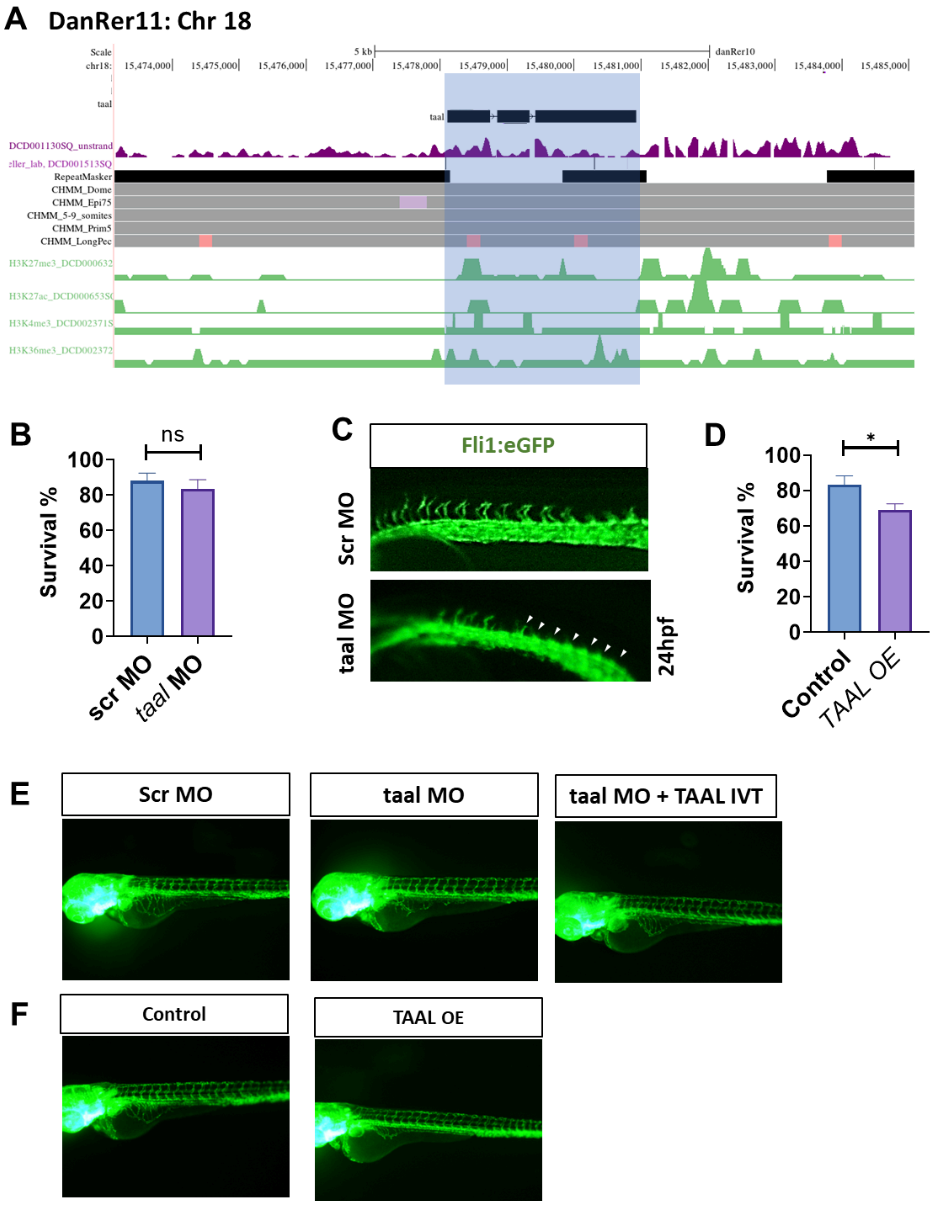


**Supplementary Figure 8:** [A] UCSC genome browser screenshot for zebrafish *taal* lncRNA locus

[B]Bar plot representing the survival of the embryos upon knockdown using *taal* morpholino. ns- not significant (two-tailed unpaired t-test)

[C]Representative images of trunk vasculature of control and morpholino-mediated *taal* knockdown Tg (fli1:eGFP) zebrafish at 24 hpf.

[D]Bar plot representing the survival of the embryos upon knockdown using *taal* morpholino. ns- not significant (two-tailed unpaired t-test)

[E-F]Raw images of angiogenic sprouting in control and Overexpressed *TAAL* injected at subintestinal vessels (SIVs) in Tg (fli1:eGFP) zebrafish.

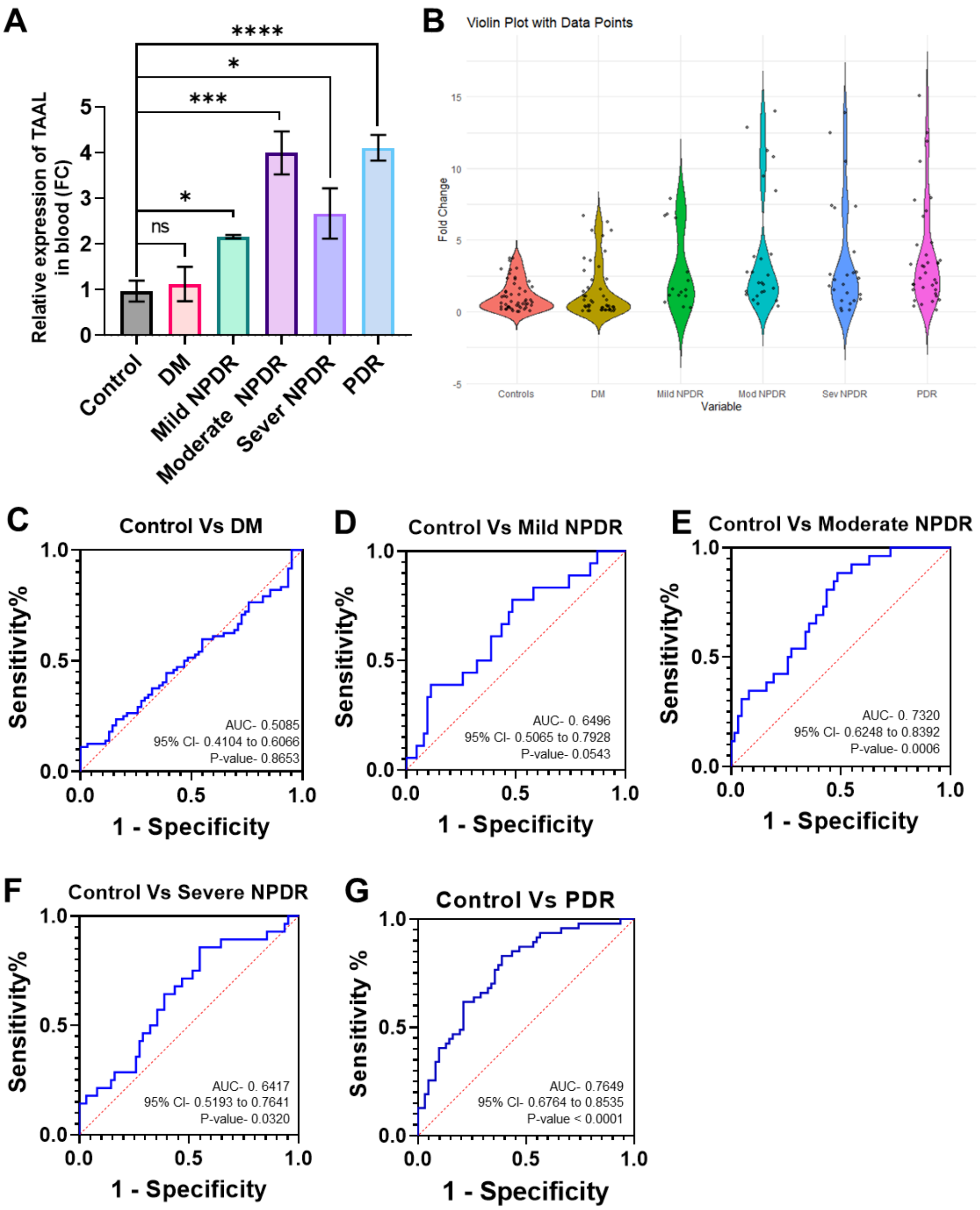


**Supplementary Figure 9:**

[A] Bar graph and violin plot with data point represent relative fold change expression of lncRNA *TAAL* in blood samples of control, diabetic mellitus (DM), Mild, Moderate and Severe nonproliferative diabetic retinopathy (NPDR) and proliferative diabetic retinopathy (PDR) patients.

[C-G] ROC curve analysis determines the sensitivity and specificity of lncRNA *TAAL* as a diagnostic biomarker for DM, mild, moderate and severe NPDR and PDR patients with vascular dysfunction compared to the control group.
